# Reciprocal regulation of p21 and Chk1 controls the Cyclin D1-RB pathway to mediate senescence onset after DNA damage-induced G2 arrest

**DOI:** 10.1101/2021.08.17.452482

**Authors:** Gerald Lossaint, Anđela Horvat, Véronique Gire, Katarina Bacevic, Karim Mrouj, Fabienne Charrier-Savournin, Virginie Georget, Daniel Fisher, Vjekoslav Dulic

**Affiliations:** Institut de Génétique Moléculaire de Montpellier (IGMM), University of Montpellier, CNRS, Montpellier, France; Institut du cancer de Montpellier, Montpellier, France; Ruđer Bošković Institute, 10000 Zagreb, Croatia; Centre de recherche en biologie cellulaire de Montpellier, (CRBM) University of Montpellier, CNRS, Montpellier, France; Institute for Stem Biology and Regenerative Medecine, Stanford University School of Medicine, Stanford, CA 94305, USA; PerkinElmer, Inc., Parc Marcel Boiteux, BP 84175 30200 Codolet, France; MRI-CRBM, BioCampus, Univ. Montpellier, CNRS, Montpellier, France

**Author notes:** These authors contributed equally to this work (Gerald Lossaint, Anđela Horvat).

## Abstract

Senescence is an irreversible proliferation withdrawal that can be initiated after DNA damage-induced cell cycle arrest in G2 phase to prevent genomic instability. Senescence onset in G2 is not well understood; it requires p53 and RB family tumour suppressors, but how they are regulated to convert a temporary cell cycle arrest into a permanent one remains unknown. Here, we show that a previously unrecognised balance between the CDK inhibitor p21 and Chk1 controls D-type cyclin-CDK activity during G2 arrest. In non-transformed cells, p21 activates RB in G2 by inhibiting Cyclin D1-CDK2/CDK4. The resulting G2 exit, which precedes appearance of senescence markers, is associated with a mitotic bypass, Chk1 inhibition and DNA damage foci reduction. In p53/RB-proficient cancer cells, compromised G2 exit correlates with sustained Chk1 activity, delayed p21 induction, untimely Cyclin E1 re-expression and genome reduplication. Chk1 depletion promotes cell cycle exit by inducing p21 binding to Cyclin D1 and Cyclin E1-CDK complexes and down-regulating CDK6, whereas Chk2 knockdown promotes RB phosphorylation and delays G2 exit. In conclusion, p21 and Chk2 oppose Chk1 to maintain RB activity, thus controlling DNA damage-induced senescence onset in G2.

## INTRODUCTION

Cellular senescence is a permanent proliferation withdrawal that can be induced by diverse stimuli such as dysfunctional telomeres, DNA damage, excessive mitogenic signalling and oncogene activation. In addition to preventing neoplastic transformation, senescence plays an important role in a number of physiological and pathological processes, and it contributes to age-related disorders and cancer (He & Sharpless, 2017). Senescent cells are characterized by large nuclei, cell hypertrophy due to intense metabolic activity, increased b-galactosidase (SA-b-gal) activity and enhanced secretion of proinflammatory molecules, known as the senescence-associated secretory phenotype (SASP; (Sharpless & Sherr, 2015)).

These routinely used markers of senescence are preceded by an irreversible DNA damage-induced cell cycle arrest that requires functional p53 and retinoblastoma protein (RB)-family tumour suppressors (Sharpless & Sherr, 2015). In replicative senescence, G1 arrest is mediated by the p53-induced CDK inhibitor p21^Waf1/Cip1^ (p21) that inhibits G1 cyclin (CycD1 and CycE1)-bound CDKs, thereby blocking inactivating phosphorylation of RB family pocket proteins and DNA replication (Dulic et al, 2000; Stein et al, 1999). This stable G1 arrest does not require the CDK4/CDK6-specific inhibitor p16^INK4A^ (p16) that accumulates only in late stages of senescence and ensures its irreversibility by preventing formation of CycD1-CDK4/CDK6 complexes (Alcorta et al, 1996; He & Sharpless, 2017; Ito et al, 2018; Sharpless & Sherr, 2015; Stein et al, 1999).

The senescence can be also triggered in G2 phase of the cell cycle (Gire & Dulic, 2015; Shaltiel et al, 2015). DNA damage-induced G2 arrest is initiated by ATM/ATR–mediated phosphorylation of checkpoint kinases Chk1 and Chk2 that, by inhibiting CDC25 phosphatases, block activation of the mitosis inducer CDK1 (Chen & Poon, 2008). Converting a temporary G2 arrest into a permanent cell cycle arrest, termed G2 exit, which precedes apparition of senescent markers, requires p21. In addition to inhibiting CycA-CDK1/2 (Bacevic et al, 2017a; Baus et al, 2003; Lossaint et al, 2011), thereby preventing CycB1-CDK1 activation (Lemmens & Lindqvist, 2019), p21 sequesters inactive CycB1-CDK1 in the nucleus (Charrier-Savournin et al, 2004; Krenning et al, 2014). This leads to downregulation of CycB1 and other mitotic regulators (Charrier-Savournin et al, 2004; Lossaint et al, 2011) *via* APC/C^Cdh1^-mediated degradation (Shaltiel et al, 2015). The G2 exit, which probably coincides with senescence onset in G2, is preceded by mitotic bypass, giving rise to stably arrested tetraploid G1 cells (Gire & Dulic, 2015; Johmura et al, 2014; Krenning et al, 2014). Additionally, p21 inhibits phosphorylation of RB family pocket proteins (Baus et al, 2003; Johmura et al, 2014) that leads to repression of E2F-dependent G2/M regulators (Jackson et al, 2005; Johmura et al, 2014). However, the identity of the RB kinase(s) that are targeted by p21 to trigger G2 exit has not been established. While CycD1-associated CDK4 and CDK6 promote G1/S progression by phosphorylating and inactivating RB (Chung et al, 2019; Topacio et al, 2019), the role of the CycD1-RB module after the restriction (R) point remains under- studied. Unlike CycE1, CycD1 is also expressed in late cell cycle phases (Chassot et al, 2008; Gookin et al, 2017; Hitomi & Stacey, 1999; Matsushime et al, 1994; Yang et al, 2006). It is therefore possible that, like at G1/S transition (Lundberg & Weinberg, 1998), CycD1-CDKs “prime” RB enabling further phosphorylation by CDK2 and/or CDK1. This is consistent with a CycD1 role as a sensor/effector of anti-proliferative cues even beyond G1/S transition (Min et al, 2020; Yang et al, 2017b). However, whether the CycD1-RB module senses DNA damage in G2 is not known.

Permanent G2 exit was proposed to serve as a safeguard mechanism preventing adaptation to the G2/M checkpoint (Baus et al, 2003), *i.e.* passage into mitosis of cells with damaged DNA, which can occur in cells lacking sufficient Chk1 activity (Feringa et al, 2018; Shaltiel et al, 2015). G2/M checkpoint adaptation also occurs in the absence of p53 or p21, and subsequent cytokinesis failure can lead to accumulation of polyploid nuclei or cell death (Bunz et al, 1998; Johmura et al, 2014). Alternatively, in p53-deficient cells experiencing persistent telomere dysfunction, sustained Chk1 and Chk2 activity coincided with prolonged G2 arrest entailing mitotic bypass and genome reduplication (Davoli et al, 2010), generating genomic instability (Davoli & de Lange, 2012). These results imply that, like p21, Chk1/Chk2 kinases stabilise the G2 arrest following continuous DNA damage, and suggest a certain redundancy between the two pathways. They also show that stable G2 arrest is not equivalent with, and does not necessarily lead to G2 exit. Paradoxically, although recent work implicated ATR/Chk1 in the senescence onset in G2 (Feringa et al, 2018; Johmura et al, 2016), DNA damage-induced senescence or terminal differentiation are all invariably associated with Chk1 downregulation (Gabai et al, 2008; Gottifredi et al, 2001; Lossaint et al, 2011; Park et al, 2015; Ullah et al, 2011). It is currently unclear whether Chk1 signalling shutdown is required for the permanent cell cycle arrest or is merely a consequence.

In this study, we investigated the kinase network that controls the transition between DNA damage-induced G2 arrest and permanent G2 exit preceding senescence. We found that in non-transformed cells the G2 exit is driven by p21-dependent inhibition of CycD1-CDK complexes that blocks RB phosphorylation and coincides with Chk1 but not Chk2 downregulation. Furthermore, sustained Chk1 activation in cancer cells is associated with impaired G2 exit and endoreplication, whereas its acute depletion strongly accelerated permanent cell cycle exit *via* p21-mediated inhibition of RB kinases. Our results suggest that, due to opposing regulation of RB phosphorylation, Chk1 inhibits, whereas p21 and Chk2 promote, the onset of senescence in G2. Finally, we uncover CDK6 downregulation as an important component of the CycD1-RB pathway control during DNA damage-induced cell cycle exit in cancer cells.

## RESULTS

### p21 targets CycD1 to inhibit RB phosphorylation and promote senescence onset after G2 arrest

RB and p21 play key roles both in senescence (Chicas et al, 2010) and in DNA damage-induced permanent cell cycle arrest in G2 (Baus et al, 2003; Johmura et al, 2014). As such, the p21- mediated inhibition of RB kinases might be a key event triggering the switch between temporary and permanent G2 arrest. Since in human diploid fibroblasts (HDF) CycD1 is expressed beyond G1/S transition (Fig. S1A; (Chassot et al, 2008)) we surmised that CycD1-RB module could serve as a sensor of anti-proliferative cues even in late cell cycle phases. We therefore investigated whether p21 inhibits RB phosphorylation and promotes DNA damage-induced senescence onset in G2 by targeting CycD1-CDK complexes.

To test this hypothesis, we exposed HDF to genotoxic drugs, previously shown to induce irreversible G2 arrest (hereafter G2 exit): ICRF-193, a DNA topoisomerase II inhibitor, which generates double strand DNA breaks in G2 phase, or bleomycin, a radiomimetic drug that causes G1 or G2 arrest (Fig. S1B, (Baus et al, 2003)). After 48h, both drugs inhibited RB phosphorylation, down-regulated CycA and Ki-67, and induced accumulation of p21 and hypo-phosphorylated p130 (Fig. 1A and Figs. S1C-E). These hallmarks of permanent cell cycle arrest (cell cycle exit), observed also upon irradiation and in replicative senescence (Fig. 1A and Fig. S1E), invariably preceded later induction of senescent markers such as increase of nuclear size (Fig. 1B), upregulation of p16^Ink4a^ and acquisition of SA-b-gal staining (Figs. S1E-G). In addition, both G1 and G2 cell cycle arrests led to accumulation of CycE1 and CycD1, a phenotype also observed in senescent cells (Fig. 1A and Fig. S1E; Dulic et al, 1993). Whereas CycE1 accumulated only after CycA degradation in p21-bound complexes, as a result of mitotic bypass (Fig. 1A and Fig. S2A), single cell immunofluorescence analysis showed that CycD1 increased already in G2-arrested cells, as documented by co-expression with CycA and CycB1 (Fig. 1C,D and Figs. S2B,C).

**Figure 1.**
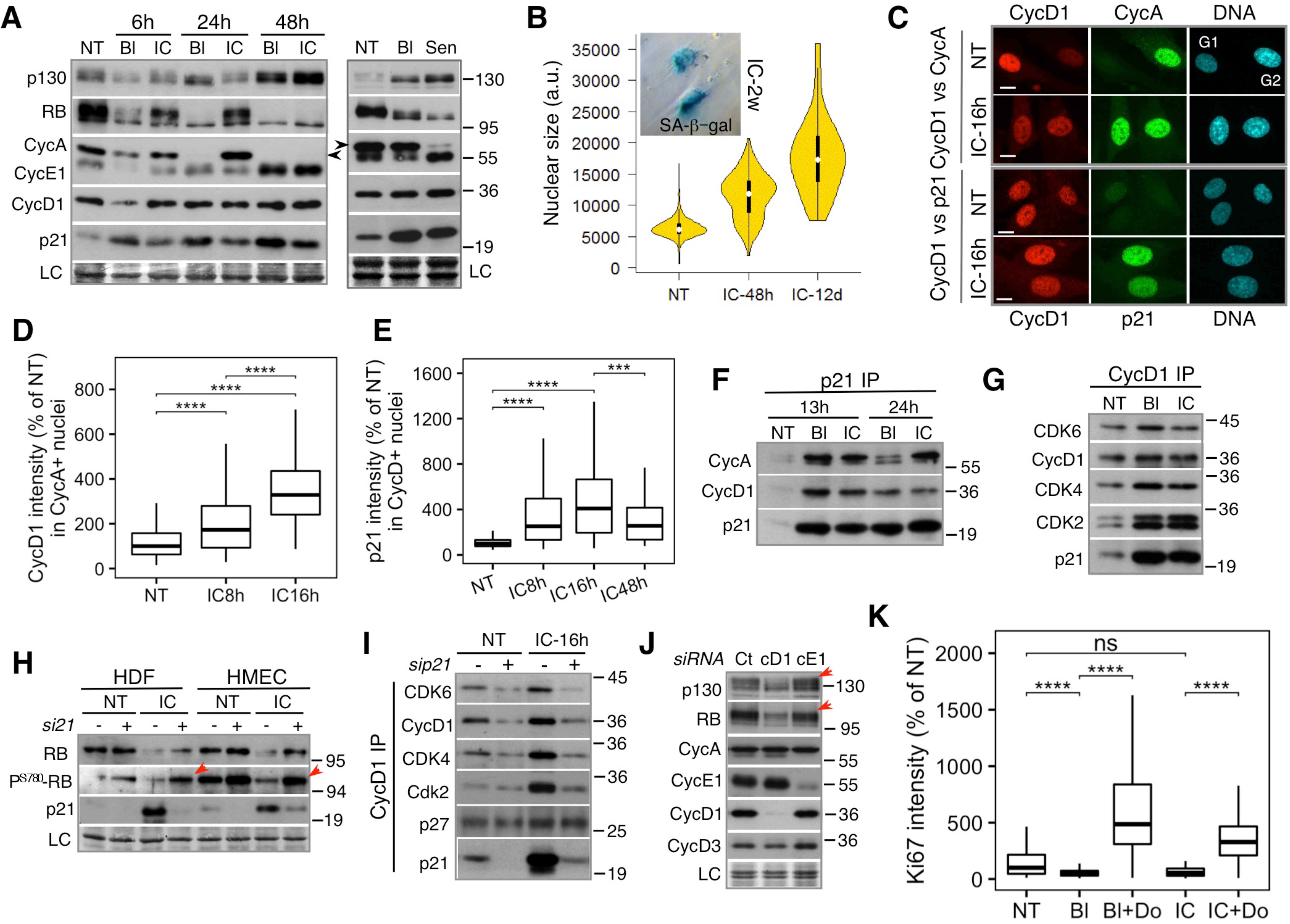
p21 targets Cyclin D1 to inhibit RB phosphorylation and promote senescence onset after G2 arrest. **A.** Immunoblots showing G1 cyclins, p21 and RB/p130 phosphorylation after DNA damage- induced cell cycle exit (left panel) and in senescent cells (right panel). HDF were exposed to bleomycin (Bl) or ICRF-193 (IC) for indicated times and for 12 hr in right panel. Sen: population doubling (PD) 74. NT, non-treated cells. Arrow shows CycA band. LC, loading control. **B.** Violin plots showing nuclear size in non-treated (NT) and HDF exposed to ICRF for 48 hr and 12 days. More than 200 cells were analysed in each experiment (n=2). Insert: phase contrast images showing β-galactosidase staining of HDF exposed to ICRF-193 (IC) for two weeks. **C.** Representative immunofluorescence images (n=3) showing co-expression of CycD1 and CycA (upper panel) or p21 (lower panel) in non-treated (NT) and HDF and exposed to ICRF- 193 (IC) for 16 hours. Bar, 10 μM. **D.** Quantification of Cyclin D1 intensity in Cyclin A-positive nuclei in non-treated (NT) and HDF exposed to ICRF-193 (IC) for 8 or 16 hours (% of NT). Pooled cells (>200) from three independent experiments. Box plot whiskers indicate 10-90% boundary. **E.** Quantification of p21 intensity in Cyclin D1-positive nuclei in non-treated (NT) and HDF exposed to ICRF-193 (IC) for indicated times (% of NT). Pooled cells (>200) from three independent experiments. Box plot whiskers indicate 10-90 % boundary. **F.** Immunoblots of the indicated proteins in p21 immunoprecipitates (IP) from extracts of non-treated (NT) and HDF exposed to bleomycin (Bl) or ICRF-193 (IC) for indicated times. **G.** Immunoblots of the indicated proteins in CycD1 immunoprecipitates (IP) from extracts of non-treated (NT) and HDF exposed to bleomycin (Bl) or ICRF-193 (IC) for 16 hr. **H.** Immunoblots showing effects of p21 KD on Cyclin D1-specific RB phosphorylation (P^S780^) in extracts from non-treated (NT) or HDF and HMEC exposed to ICRF-193 (IC) for 16 hr. Arrows shows increased P^S780^-RB after p21 KD (+). LC, loading control. **I.** Immunoblots of the indicated proteins in CycD1 immunoprecipitates (IP) from extracts of non-treated (NT) or HDF exposed to ICRF-193 (IC) for 16 hr that were previously depleted (+) or not (-) for p21 (sip21). **J.** Immunoblots showing effects of CycD1 and CycE1 knockdown on p130 and RB phosphorylation (arrows). HDF were depleted for CycD1 (cD1) or CycE1 (cE1) for 24 hr. Ct, siRNA control. LC, loading control. **K.** Quantification of Ki67 intensity in T_121_-expressing fibroblasts exposed to bleomycin (Bl) or ICRF-193 (IC) for 48 hr (% of NT). Doxycycline (Do) was added 12 hr before exposure to genotoxic agents. Pooled cells (>100) from two independent experiments. Box plot whiskers indicate 10-90 % boundary. NT, non-treated cells. Loading controls (LC) were amido-black stained membranes. Data are representative or mean +/- SEM of at least two independent experiments. *P* values were calculated with two-sided Student’s *t* test; ***P ≤ 0.001, ****P ≤ 0.0001.

Next, we asked whether p21 targets CycD1-CDK complexes in G2 arrested cells. As shown by immunofluorescence (Fig. 1C, E and Figs. S2D-F) and immunoblots of p21 and CycD1 immunoprecipitates (Fig. 1F,G), G2 arrest correlated with increasing co-expression and binding of both CycD1 and CycA with p21 (Fig. 1F). In addition to CycD1-CDK4, arrested cells also accumulated CycD1-CDK2 complexes (Fig. 1G), which have previously been observed in senescent fibroblasts (Dulic et al, 1993; Stein et al, 1999). These data imply that p21 simultaneously binds CycA-CDK1/2 that drives mitosis, and CycD1-CDK2/4/6, that phosphorylate RB. Thus, preventing p21 induction might stimulate CycD1-dependent RB phosphorylation after G2 arrest. Indeed, CycD1-specific RB phosphorylation (P^S780^) was promoted by p21 knock-down (KD; Fig 1H) or expression of HPV16-E6 oncoprotein, which degrades p53 and prevents p21 induction (Baus et al, 2003) compromising the cell cycle exit (Figs. S3A-C). Moreover, p21 KD abolished accumulation of CycD1-CDK2/4/6 complexes in G2-arrested cells (Fig. 1I), linking their stabilization by p21 to G2 exit.

The above results suggested that CycD1-CDK complexes might also control RB activity beyond the G1/S transition. Therefore, we knocked down CycD1 to assess their contribution to pocket protein phosphorylation in proliferating cells. CycD1 KD strongly reduced RB and p130 phosphorylation despite unchanged CycD3, CycE1 and CycA levels (Fig. 1J) showing that these cyclins cannot fully compensate for CycD1 loss. Indeed, CycE1 depletion less affected RB and p130 phosphorylation (Fig. 1J). Finally, to validate the key role of RB as p21 target in inducing the G2 exit, we conditionally expressed SV40 large T antigen mutant (T_121_), which specifically inactivates RB orthologs after addition of doxycycline (Conklin et al, 2012). RB family inactivation abolished the cell cycle arrest and maintained Ki-67 and CycA expression upon exposure to ICRF- 193 or bleomycin despite high p21 levels (Fig. 1K and Figs. S3D,E). Collectively, these results suggest that p21 initiates G2 cell cycle exit and subsequent senescence onset by binding to and inhibiting CycD1-CDK complexes, thus blocking RB phosphorylation (Fig. S3F).

### G2 exit is preceded by p53-dependent suppression of Chk1 signalling

Next, we asked whether p21 is sufficient to ensure stable G2 arrest preceding G2 exit or it acts in synergy with Chk1/Chk2. The robustness of the G2/M checkpoint depends on the level of DNA damage above a certain threshold (Lobrich & Jeggo, 2007). Therefore, conversion of G2 arrest to permanent G2 exit might depend on the strength of DNA damage signalling, with higher levels translating into increased Chk1/Chk2 activity and p21 induction. To test this hypothesis, HDF were synchronized in early S-phase (favouring G2 arrest) and exposed to two different γ-ray doses (5 and 10 Gy; Fig. S4A). Both doses rapidly upregulated p21 and induced cell cycle exit, as documented by the absence of RB phosphorylation and CycA expression and stabilisation of G1 cyclins (Fig. 2A). However, whereas after 10 Gy cells arrested predominantly in G2, at 5 Gy most cells entered mitosis and arrested in G1 (Fig. 2B, and Fig. S4A). Accordingly, robust G2 arrest correlated with initially stronger Chk1 phosphorylation (Fig. 2A, 10 Gy). In contrast, p21 induction and Chk2 phosphorylation were comparable between the two γ-ray doses (Fig. 2A). This suggests that the intensity of Chk1 activation, rather than p21 upregulation, determines whether cells arrest and exit the cell cycle in G2, or progress into mitosis and exit in G1. Similar conclusions regarding the role of Chk1 in stabilizing G2 exit were obtained with another cellular model (Johmura et al, 2016). If this is true, then stronger Chk1 activation should correlate with more robust G2 arrest even if cells cannot induce p21. Indeed, whereas at 10 Gy most E6-expressing cells arrested in G2, at 5 Gy they progressed into the next cell cycle (Fig. 2C and Fig. S4B). Likewise, time-lapse studies showed that stronger Chk1 activation by bleomycin correlated with a robust G2 arrest, whereas ICRF-193-treated HDF-E6 exhibiting weaker Chk1 phosphorylation, entered mitosis despite strong Chk2 activation (Fig. 2E,F and Fig. S4C). However, both G1- and G2-arrested HDF-E6 cells failed to exit the cell cycle, as shown by persistent RB hyper-phosphorylation (Fig. 2D).

**Figure 2.**
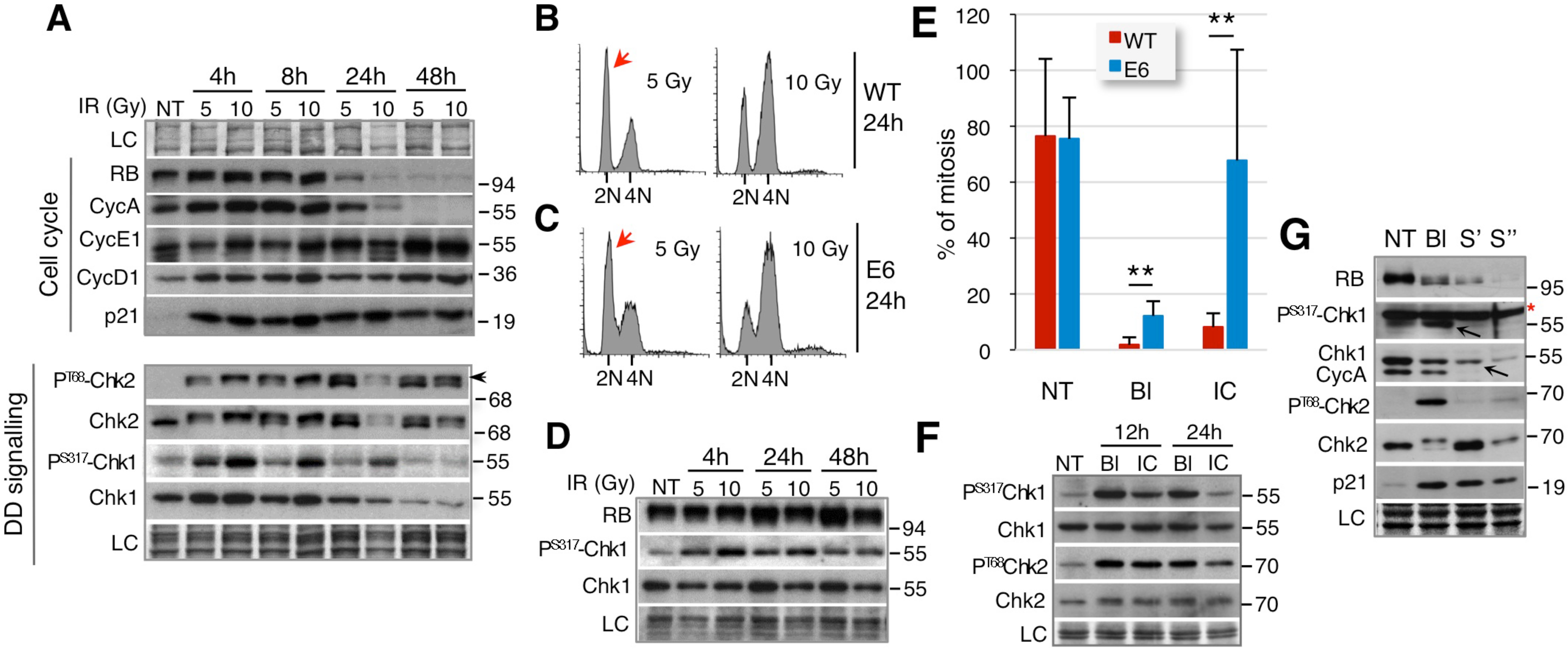
G2 exit is preceded by p53-dependent suppression of Chk1 signalling. **A.** Immunoblots of the indicated cell cycle regulators (upper panel) and DNA damage signalling proteins (lower panel) in extracts from synchronized HDF exposed to ionizing (γ) radiation (IR; 5 or 10 Gy) 16 hours after release from G0. NT, non-treated cells; LC, loading control. Arrows: phosphorylation-induced SDS-PAGE mobility shift of Chk2. **B.** Flow cytometry DNA content profiles of irradiated wild-type (WT) and (**C**) HPV16-E6 - expressing HDF (E6) after 24 hr. Red arrows: G1 cells. **D.** Immunoblots showing persistent RB phosphorylation and the absence of Chk1 down- regulation in irradiated (5 or 10 Gy) asynchronous HDF-E6. NT, non-treated cells; LC, loading control. (WT in Fig. S3C). **E.** Video-microscopy showing percent of cells entering mitosis in wild-type (WT) and HPV16-E6-expressing HDF exposed to ICRF-193 (IC) or bleomycin (BL) for 24 hours (mean +/- SD of 3 fields from two separate experiments). *P* values were calculated with two-sided Student’s *t* test; **P ≤ 0.01. **F.** Immunoblots showing phosphorylation of Chk1 (P^S317^) and Chk2 (P^T68^) in extracts from HDF expressing HPV16-E6 exposed to ICRF193 (IC) or bleomycin (Bl) for indicated times. NT, non-treated cells; LC, loading control (WT in Fig. S4C). **G.** Immunoblots of the indicated cell cycle regulators and DNA damage signalling proteins in extracts of early passage (population doubling (PD) 30), non-treated (NT) or exposed to bleomycin (Bl-12h), and senescent HDF (S’ - PD 74; S’’ - PD 84). Arrows: P^S317^-Chk1 and Chk1 bands; Red asterisk: nonspecific band. Loading controls (LC) are amido-black stained membranes.

Surprisingly, given its key role in G2 arrest, Chk1 phosphorylation and protein levels rapidly decreased before the onset of cell cycle exit, contrasting persistent Chk2 phosphorylation (Fig. 2A). This transient Chk1 phosphorylation was not specific to irradiation since it was also observed in cells exposed to bleomycin or ICRF-193 (Fig. S4C,D). By contrast, both Chk1 phosphorylation and protein levels were maintained in HDF-E6 arrested in G2 by radiation or bleomycin (Fig. 2D,F). Thus, prolonged Chk1 phosphorylation coincided with impaired cell cycle exit. Conversely, Chk1 was strongly downregulated in DNA damage-induced and replicative senescence (Fig. 2G and Fig. S4E). Taken together, these results highlight a correlation between G2 exit and suppression of Chk1, but not Chk2, signalling by the p53/p21 pathway suggesting their differential roles in G2 arrest/G2 exit conversion.

### G2 exit associates with gH2AX downregulation and reduction of DNA damage foci

We wondered if Chk1 downregulation might be a part of general DNA damage signalling switch-off associated with the onset of G2 exit. To explore this possibility, we analysed by immunofluorescence the gH2AX signal intensity and the number of DNA damage-induced foci at different times after exposure to ICRF-193 in both wild-type and E6-expressing HDF. Indeed, G2 exit (48-72h) correlated with a marked reduction of both number of γH2AX/53BP1 foci and γH2AX signal that were strongly attenuated by E6-expression (Fig. 3A-D). Downregulation of the DNA damage response signalling is unlikely due to DNA damage repair (Chowdhury et al, 2005) or checkpoint recovery (Macurek et al, 2010) since the experimental conditions induced massive DNA damage, causing senescence. Moreover, reduction of γH2AX/53BP1 foci correlated with an increase in size (Fig. 3A,B) that might reflect their clustering associated with delayed repair (Aymard et al, 2017). These large foci resemble 53BP1 nuclear bodies that form in the G1 phase after unrepaired DNA damage or unresolved replication stress in the preceding cycle (Lukas et al, 2011). Significantly, in addition to the absence of Chk1 phosphorylation (Fig. 2G), most senescent cells also exhibited low gH2AX signal and reduced number of larger DNA damage foci (Fig. 3E). Based on our results, we suggest that implementation of G2 exit and senescence onset involves downregulation of Chk1 and ATM/ATR signalling *via* the p53/RB pathway (Fig. 3F).

**Figure 3.**
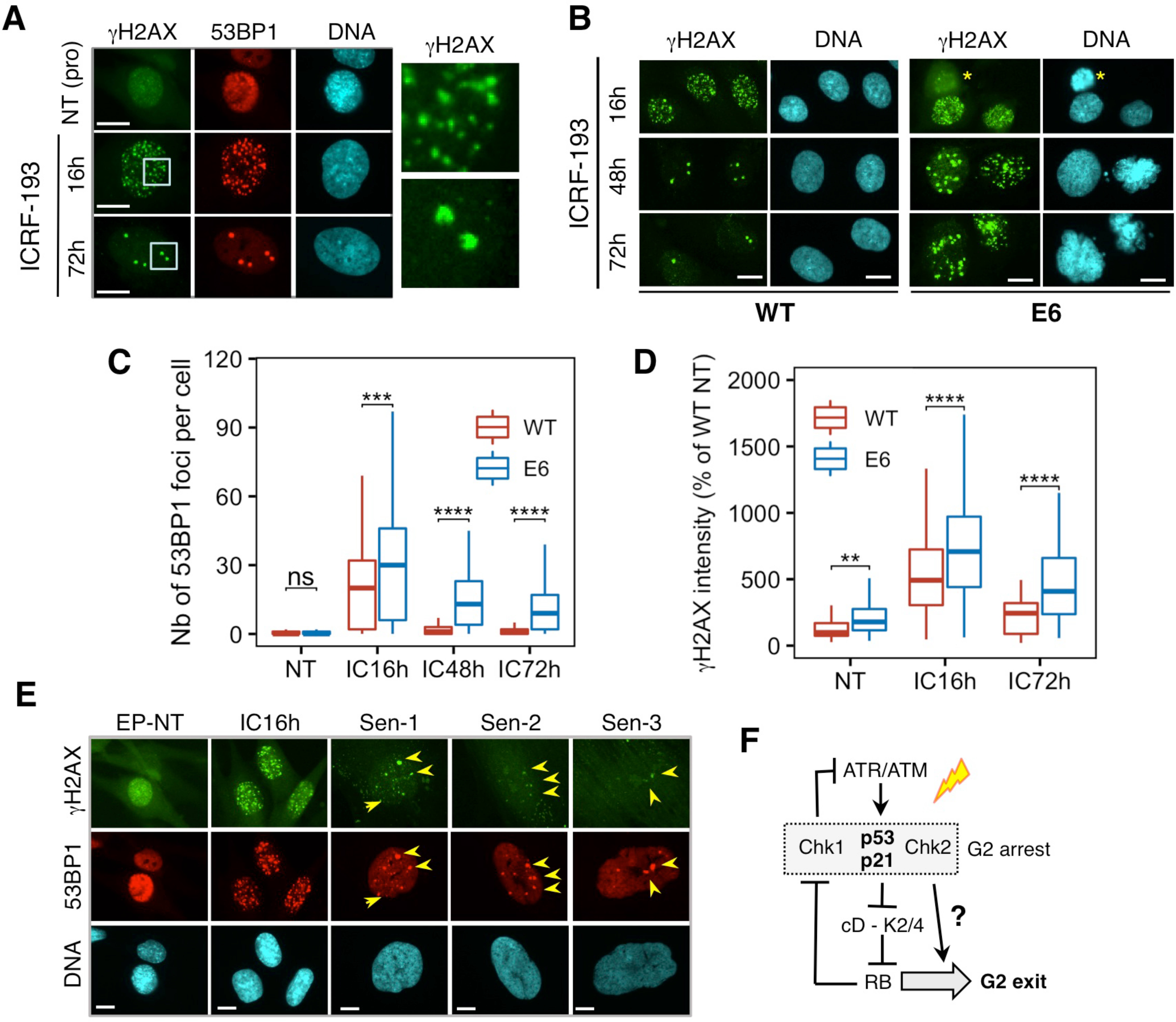
G2 exit associates with γH2AX downregulation and reduction of DNA damage foci. **A.** Representative immunofluorescence images (n=2) showing co-localization of γH2AX and 53BP1 foci in non-treated (NT, cell in prophase) and HDF exposed to ICRF-193 for indicated times. Right panel shows magnified area to appreciate differences in size/shape of gH2AX foci in cells exposed to ICRF-193 for 16 hr (up) and 72 hr (down). Bar, 10 μM. **B.** Representative immunofluorescence images (n=3) showing γH2AX foci in WT and E6- expressing HDF exposed to ICRF-193 for indicated times. Asterisk (*) denotes cell in prophase. Bar, 10 μM. **C.** Quantification of 53BP1 foci in WT and E6-expressing HDF exposed to ICRF-193 (IC) for indicated times. NT, non-treated cells. Pooled cells (>200) from three independent experiments. Box plot whiskers indicate 10-90 % boundary. **D.** Quantification of γH2AX signal intensity in the nuclei of WT and E6-expressing HDF exposed to ICRF-193 at indicated timepoints (percent of NT cells). Pooled cells (>100) from three independent experiments. Box plot whiskers indicate 10-90 % boundary. NT, non- treated cells. **E.** Immunofluorescence images showing co-expression and co-localization of γH2AX and 53BP1 foci in non-treated (NT) early passage HDF (EP-NT), HDF exposed to ICRF-193 (IC16h) and in several representative senescent HDF (PD 84). Arrows indicate γH2AX/53BP1 foci in senescent cells. Bar, 10 μM. **F.** Proposed roles for p53/p21 and CycD1-RB modules in suppressing Ckh1 and ATM/ATR signalling during G2 arrest-G2 exit switch. *P* values were calculated with two-sided Student’s *t* test; ***P ≤ 0.001, ****P ≤ 0.0001.

### Sustained Chk1 activation after G2 arrest coincides with altered mitotic bypass and delayed G2 exit in U2OS cells

If G2 exit requires Chk1 downregulation, then one might expect that prolonged Chk1 activity due to sustained DNA damage, as previously observed in p53/RB-proficient cancer cell lines (Lossaint et al, 2011) or p53-deficient cells (Davoli et al, 2010), would be incompatible with its onset. To address this question, we studied U2OS osteosarcoma cells that, when exposed to genotoxic agents, arrest predominantly in G2 due to a deficient ATM signalling and persistent Chk1 activation (Figs. S5A,B (Kleiblova et al, 2013; Lossaint et al, 2011)). Similarly, g-irradiation (IR) led to delayed p21 induction and sustained dose-dependent Chk1 phosphorylation, which correlated well with robustness of G2 arrest (Fig. 4A, B, upper panel).

**Figure 4.**
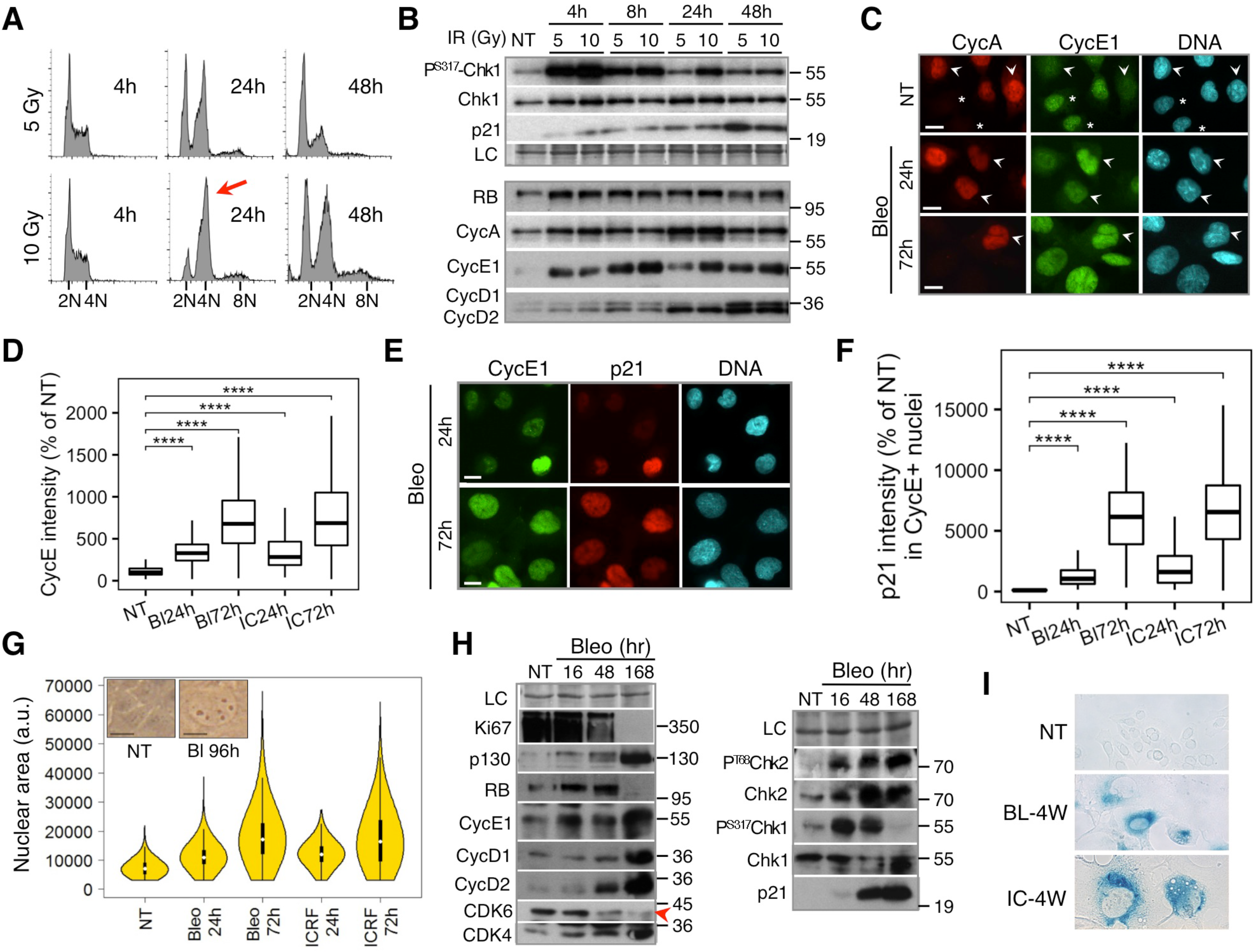
Sustained Chk1 activation after G2 arrest coincides with altered mitotic bypass and delayed G2 exit in U2OS cells. **A.** Flow cytometry DNA content profiles of U2OS cells at indicated times after γ irradiation (5 or 10 Gy). Note more efficient G2 arrest in cells irradiated with 10 Gy (red arrow). **B.** Immunoblots showing changes in Chk1 phosphorylation, p21 (upper panel) and denoted cell cycle regulators (lower panel) in U2OS cells exposed to two doses of ionizing (γ) radiation (IR; 5 or 10 Gy). NT, non-treated cells; LC, loading control. **C.** Representative immunofluorescence images (n=2) showing CycE1/CycA co-expression in non-treated (NT) and U2OS cells exposed to bleomycin (Bleo) for 24 or 72 hours. Asterisk in NT indicate G1 cells that lack CycA whereas arrowheads show CycA-positive cells with low (NT) and high (Bleo) CycE1 expression. Bar, 10 μM. **D.** Quantification of CycE1 intensity in U2OS exposed to bleomycin (BL) or ICRF-193 (IC) for 24 or 72 hours (% of NT). Pooled cells (>200) from two independent experiments. Box plot whiskers indicate 10-90 % boundary. NT, non-treated cells. **E.** Representative immunofluorescence images (n=2) showing CycE1/p21 co-expression in U2OS cells exposed to bleomycin (Bleo) for indicated times. Bar, 10 μM. **F.** Quantification of p21 intensity in CycE1-expressing U2OS cells exposed to bleomycin for indicated times (% of NT). Pooled cells (>200) from two independent experiments. Box plot whiskers indicate 10-90 % boundary. NT, non-treated cells. **G.** Violin plots showing nuclear size in non-treated (NT) and U2OS exposed for 24 and 72 hours to bleomycin (Bleo) or ICRF-193 (ICRF). More than 500 cells were analyzed in each experiment (n=2). Inserts are micrographs showing NT and cells exposed for 96 hr to bleomycin. Bar, 10 μM. **H.** Immunoblots showing effects of prolonged exposure to bleomycin (Bleo) on indicated cell cycle regulators (left panel) and DNA damage signalling effectors (right panel) in U2OS cells. NT, non-treated cells; LC, loading control. Arrowhead shows CDK6 downregulation. **I.** Phase contrast images showing β-galactosidase staining of U2OS cells exposed to bleomycin (Bl) or ICRF-193 (IC) for 4 weeks. Loading controls (LC) are amido-black stained membranes. *P* values were calculated with two-sided Student’s *t* test; **** P ≤ 0.0001.

Consistent with our hypothesis, cell cycle exit was strongly impaired in G2-arrested U2OS cells, as documented by continuous RB hyperphosphorylation and CycA expression in both irradiated and cells exposed to genotoxic agents (Fig. 4B-lower panel and Figs. S5B-D). In addition, immunofluorescence results showed that re-accumulation of CycE1 occurred before CycA downregulation revealing altered mitotic bypass that was associated with DNA re-replication (Fig. 4B,C and Figs. S6A-D). However, extended G2 arrest entailed further upregulation of G1 cyclins that was coupled with reduced CycA and increased co-expression of p21 (Fig. 4B-F and Figs. S6A,B,E-H). Concomitant decrease of CycA-positive cells (Fig 4C and Figs. S5C,D and S6A,E) and nuclear size augmentation (Fig. 4C,E,G) indicated ongoing G2 arrest/G2 exit conversion. Indeed, prolonged exposure to bleomycin further upregulated p21 and led to permanent cell cycle exit, as shown by complete loss of Ki67, the absence of RB phosphorylation, strong accumulation of hypo-phosphorylated p130 and G1 cyclins and marked CDK6 downregulation (Fig. 4H-left). Notably, G2 exit coincided with a suppression of Chk1 signalling, but increased phosphorylation of Chk2 (Fig. 4H-right), further highlighting differences in regulation between the two kinases. Moreover, G2 exit was followed by senescence, as documented by SA-β-gal staining, which became apparent only after 2-week exposure to genotoxic agents (Fig. 4I and Fig. S6I).

Thus, sustained Chk1 activity in G2-arrested U2OS cells associates with delayed p21 induction and impaired cell cycle exit, while G2 exit and senescence onset correlate with downregulated Chk1, persistent Chk2 activity and strong accumulation of G1 cyclins.

### Acute Chk1, but not Chk2, depletion promotes DNA damage-induced cell cycle exit by inhibiting RB kinases

The striking difference in behaviour between Chk1 and Chk2 suggested that these two kinases play opposing roles at the onset of G2 exit. Based on our results and those showing that Chk1 inhibits p21 expression (Beckerman et al, 2009; Hsu et al, 2019) while Chk2 inhibits CDK (Chen & Poon, 2008), we predicted that the absence of Chk1 should promote DNA damage-induced cell cycle exit, whereas reduced Chk2 would delay it (Fig. 5A). To test this hypothesis, we knocked down each kinase in U2OS cells, which did not significantly affect cell proliferation (Fig. S7A), before exposure to genotoxic drugs. In agreement with previous work (Bacevic et al, 2017a; Lossaint et al, 2011), Chk1 KD, but not Chk2 KD, abrogated G2 arrest by bleomycin, although, based on FACS results, this was not clear for ICRF-193 (Fig. 5B). Further inspection by video- microscopy showed that the persistence of cells with 4N DNA content upon Chk1 KD in the presence of ICRF-193 was due to accumulation of bi-nucleate G1 cells generated after cytokinesis failure (Fig. 5C).

**Figure 5.**
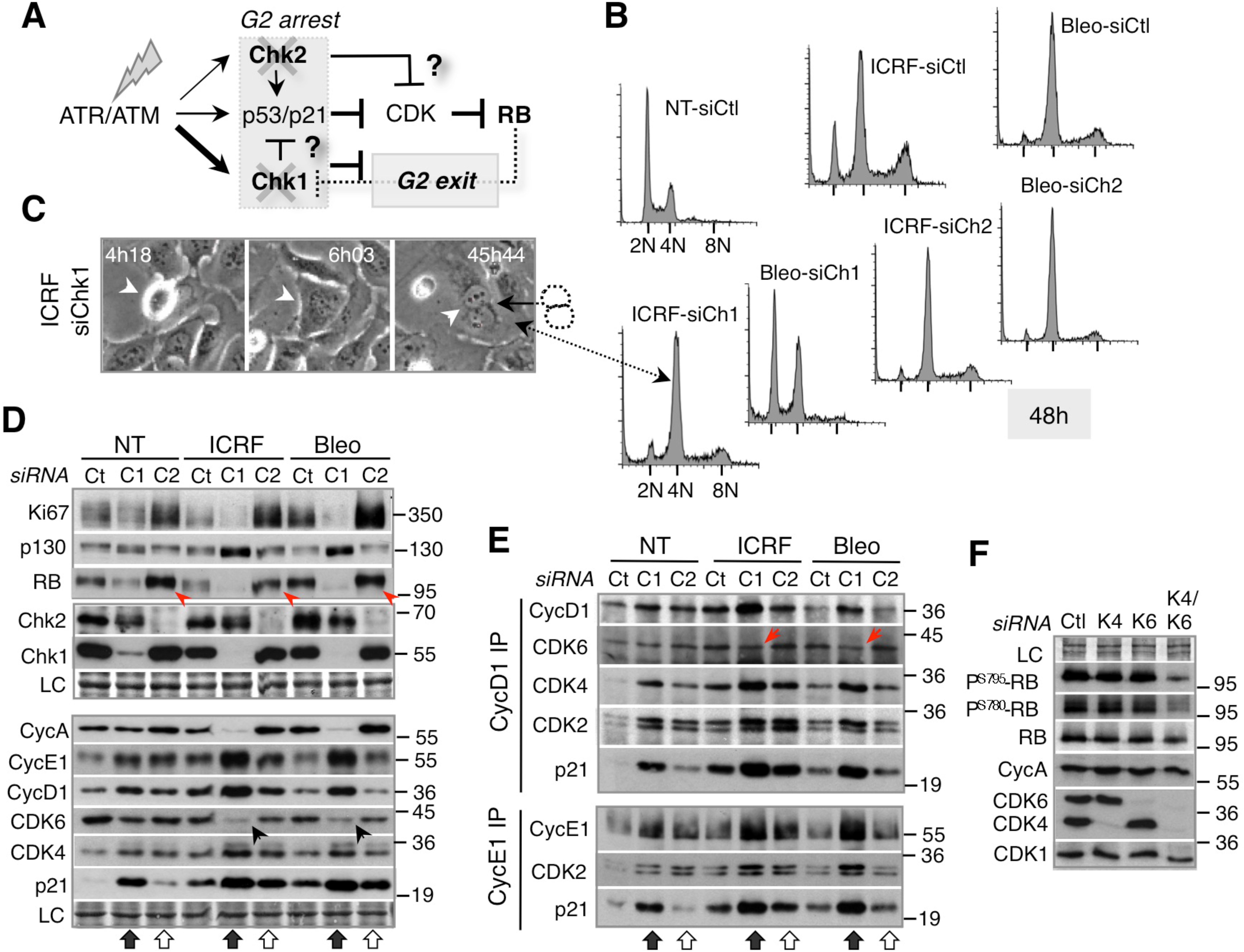
Acute Chk1, but not Chk2, depletion promotes DNA damage-induced cell cycle exit by inhibiting RB kinases. **A.** Model: Expected effects of Chk1 and Chk2 knockdown on RB pathway and DNA damage-induced G2 arrest/exit switch. **B.** Flow cytometry DNA content profiles of non-treated (NT-siCtl) and siCtl, siChk1 and siChk2 U2OS cells exposed to bleomycin (Bleo) and ICRF-193 (ICRF) for 48 hr. See Fig.5SA for NT-siChk1 and NT-siChk2. **C.** Phase contrast images from video-microscopy sequences showing binuclear daughter cell after mitosis and cytokinesis failure (arrowheads) in ICRF- treated U2OS Chk1 KD cells. Time after the addition of the drug is indicated. **D.** Immunoblots showing effects of Chk1 (siCh1-black arrow) and Ckh2 (siCh2-white arrow) knockdown on Ki67 and the indicated cell cycle regulators in non-treated (NT) and U2OS cells exposed to ICRF-193 (ICRF) or bleomycin (Bleo) for 48 hr. LC, loading control. See Fig. S5B for 16 hr time-point. **E.** Immunoblot analysis of Cyclin D1 and Cyclin E1 immunoprecipitates (IP) showing effects of Chk1 (Ch1-black arrow) and Chk2 (Ch2-white arrow) knockdown on cyclin-CDK-p21 complexes in U2OS exposed to ICRF-193 (ICRF) and bleomycin (Bleo) for 48 hr. Arrows: reduction of CDK6 in CycD1 complexes. **F.** Immunoblots showing effects of CDK4 (K4), CDK6 (K6) or double CDK4 / CDK6 (K4/K6) knockdown on Cyclin D1-specific RB phosphorylation (P^S795^, P^S780^) in U2OS cells. Loading controls (LC) are amido-black stained membranes.

As predicted, Chk1 depletion potently accelerated cell cycle exit upon DNA damage (detectable after 16h), as shown by downregulation of Ki67 and CycA, rapid inhibition of RB phosphorylation and accumulation of hypo-phosphorylated p130 and G1 cyclins (Fig 5D and Fig. S7B). Consistent with these results, Chk1 KD upregulated p21, reduced CDK6 (Fig. 5D and Fig. S7B) and strongly increased accumulation of p21-bound CycD1-CDK2/4 and CycE1-CDK2 complexes (Fig. 5E). This suggests that Chk1 KD accelerates cell cycle exit by p21-dependent inhibition of RB kinases and CDK6 downregulation. Strikingly, Chk2 KD produced exactly the opposite effect: increased Ki67, RB phosphorylation and CycA levels, and it did not upregulate G1 cyclins or induce accumulation of p21-bound CycE1 or CycD1 complexes (Fig. 5D, E).

In addition, our results revealed a difference in regulation of CDK4 and CDK6 in U2OS cells. Whereas both prolonged DNA damage (Fig. 4H) and Chk1 depletion (Fig. 5D) reduced CDK6, CDK4 levels increased in these situations, accumulating in p21-bound CycD1 complexes (Fig. 5E). To assess the respective contributions of CDK4 and CDK6 on RB regulation, we knocked down CDK4, CDK6 or both kinases. Although neither CDK4 KD nor CDK6 KD alone significantly affected RB phosphorylation, it was strongly reduced after double CDK4/6 KD despite the presence of CycA (Fig. 5F). These results highlight the importance of CDK6 downregulation in DNA damage-induced cell cycle exit and senescence onset.

Taken together, our data suggest that Chk1 and Chk2 play opposing roles in DNA damage-induced G2 exit/senescence onset in U2OS cells (Fig. 5A). Whereas sustained Chk1 activity stabilizes G2 arrest but antagonises cell cycle exit, possibly by preventing p21-mediated inactivation of RB kinases, Chk2 might promote the latter by preventing RB phosphorylation.

### Accelerated cell cycle exit by Chk1 knockdown in cancer cells is mediated by p21 induction and CDK6 downregulation

Accelerated DNA damage-induced cell cycle exit upon Chk1 KD coincided with p21 induction and CDK6 downregulation. We therefore asked whether p21 depletion would prevent this phenotype. To this end we studied the effect of p21 KD, Chk1 KD and double Chk1/p21 KD (DKD) on cell cycle exit hallmarks in U2OS cells that were exposed to bleomycin for 48 h.

Unlike Chk1 KD, p21 KD did not abrogate G2 arrest, showing that sustained Chk1 activity can prevent mitosis even after prolonged DNA damage (Fig. 6A and Fig. S8A). In contrast, acute p21 depletion increased Ki67 levels and p130/RB phosphorylation, abolished G1 cyclin accumulation and supported Chk1 phosphorylation even after prolonged DNA damage (Fig. 6B, and Figs. S8B,C). Consistent with these results, ATM inhibition by KU-55933 (Ku), which strongly reduced p21 induction and Chk2 activation, did not abrogate G2 arrest but it both increased RB phosphorylation and stabilised Chk1 phosphorylation (Fig. 6C and Fig. S8D). In contrast, caffeine, which inhibits both ATR and ATM, abrogated G2 arrest and failed to promote RB phosphorylation (Fig. 6C and Fig. S8D). These results confirmed an antagonism between Chk1 and ATM/p21/RB-mediated G2 exit. As predicted by our model, Chk1/p21 DKD abolished DNA damage-induced cell cycle exit (Fig. 6B and Fig. S8B), which resulted in unchecked cell cycle progression, mitotic catastrophes and, ultimately, cell death (Fig. 6A,D and Figs. S8A,E). Moreover, p21 KD counteracted Chk1 KD-mediated CDK6 downregulation (Fig. 6B and Fig. S8B), presumably contributing to persistent RB phosphorylation. Thus, our results confirmed the key role of p21 in Chk1 KD-induced senescence onset in U2OS cells.

**Figure 6.**
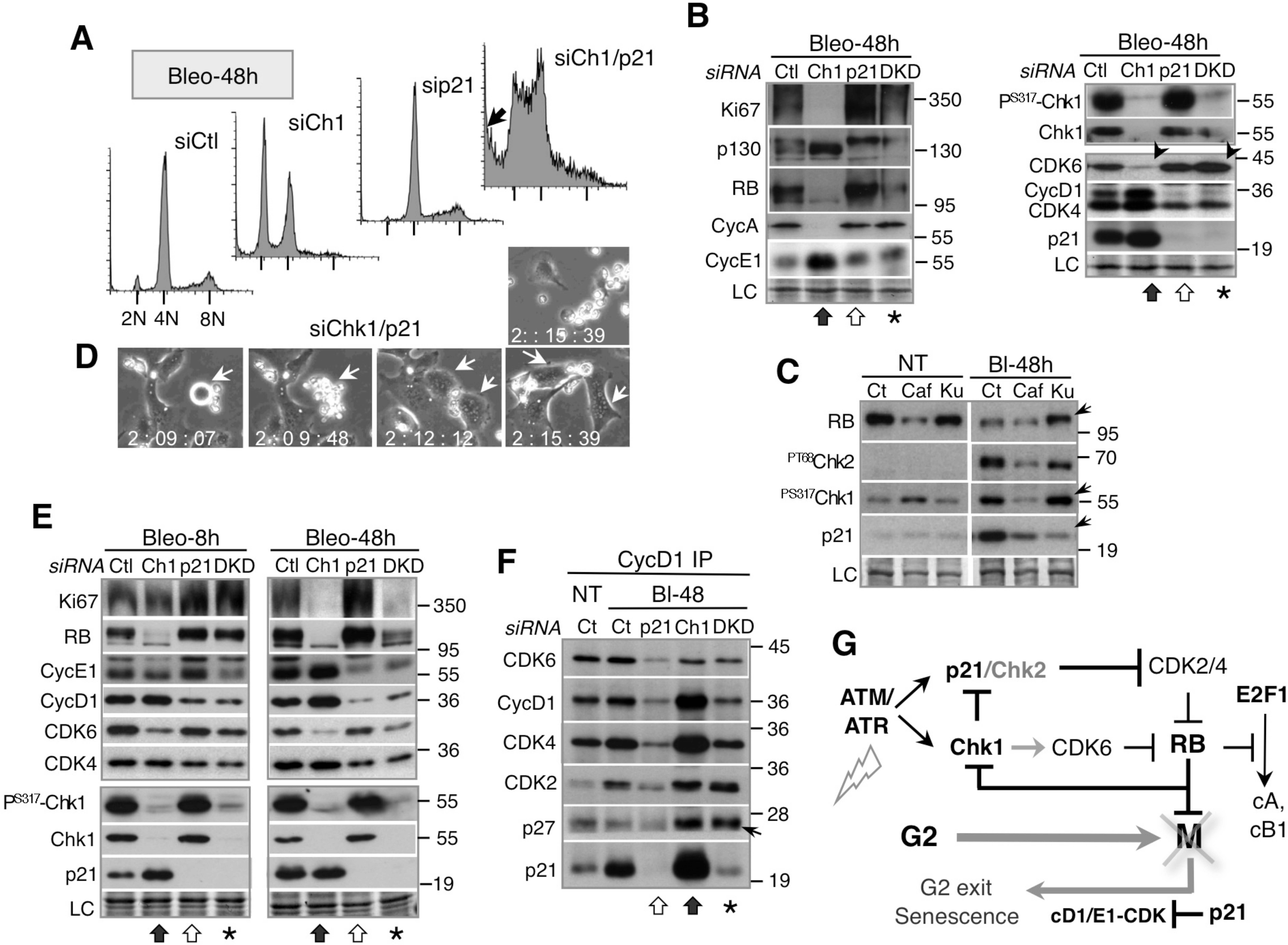
Accelerated cell cycle exit by Chk1 knockdown in cancer cells is mediated by p21 induction and CDK6 downregulation. **A.** Flow cytometry DNA content profiles of control (siCtl) and U2OS cells depleted for Chk1 (siCh1), p21 (sip21) or both (siCh1/21) proteins exposed to bleomycin (Bleo) for 48 hours. Arrow indicates sub-G1 population representing death cells. See Fig. S8A for effects of respective knockdowns on non-treated and U2OS exposed to bleomycin for 16 hours. **B.** Immunoblots showing effects of Chk1 (Ch1), p21 or double p21/Chk1 (DKD) knockdown on Ki67 and indicated cell cycle regulators and DNA damage signalling effectors in U2OS cells exposed to bleomycin for 48 hr. LC, loading control. See Fig. S8B for effects of respective knockdowns on non-treated and U2OS exposed to bleomycin for 16 hr. **C.** Immunoblots showing effects of caffeine (Caf) and ATM inhibitor KU-55933 (Ku) on RB, P^T68^-Chk2, P^S317^-Chk1 and p21 in non-treated (NT) and U2OS cells exposed to bleomycin for 48 hr (arrows). LC, loading control. **D.** Representative phase contrast images from a video-microscopy sequence showing fragmented nuclei after mitotic catastrophe in bleomycin-treated double Ckh1/p21 KD cells. Time (days, hours) after drug addition is indicated. Arrows show mitotic cell and resulting daughter cells. Most right-hand image (upper panel) shows cell debris from the same sequence. Complete field is shown in Fig. S8D. **E.** Immunoblots showing effects of Chk1 (Ch1), p21 or double p21/Chk1 (DKD) knockdowns on Ki67 and the indicated cell cycle regulators in HCT-116 cells exposed to bleomycin (Bleo) for 8 and 48 hours. LC, loading control. **F.** Immunoblots of CycD1 immunoprecipitates (IP) showing effects of Chk1 (Ch1), p21 or double p21/Chk1 (DKD) knockdowns on cyclin-CDK-CKI complexes in HCT-116 cells exposed to bleomycin (Bl) for 48 hr. Loading controls (LC) were amido-black stained membranes. Black arrow: Chk1 KD; White arrow; p21 KD; asterisk: DKD. **G.** Model: Proposed roles of Chk1, Chk2 and p21-CycD-RB axis in DNA damage-induced senescence onset after permanent G2 arrest (G2 exit). See text for details.

Next, we sought to validate the respective roles of Chk1 and p21 in the onset of G2 exit in another p53/RB-proficient cancer cell line, HCT-116. As with U2OS, RB phosphorylation and Ki67 levels persisted after bleomycin-induced G2 arrest correlating with sustained Chk1 activation (Fig. 6E and Figs. S9A,B). Chk1 KD abrogated G2 arrest and induced permanent cell cycle exit (low Ki67, unphosphorylated RB), which was associated with upregulation of p21 and G1 cyclins, strong accumulation of p21-bound CycD1-CDK2/CDK4 complexes and CDK6 downregulation (Fig. 6E,F and Fig. S9A). Conversely, while failed to abrogate G2 arrest, p21 KD increased RB phosphorylation and prevented both CycD1/CycE1 upregulation (Fig. 6E, and Figs. 9SA,B) and accumulation of CycD1-CDK complexes (Fig. 6F). Furthermore, consistent with its role in mediating Chk1 KD-induced cell cycle exit, p21 KD fully restored RB phosphorylation (Fig. 6E- 8h-DKD) and abolished upregulation of CycD1-CDK complexes (Fig. 6F-DKD). However, after prolonged DNA damage DKD only partially rescued Ki67 expression and RB phosphorylation in HCT-116 cells (Fig. 6E-48 h, asterisk). The latter might be due to upregulation of CDK inhibitor p27^Kip1^ in CycD1-CDK complexes (Fig. 6F, arrow) and reduced CDK6 levels (Fig. 6E). Moreover, like in U2OS, DKD promoted cell death by abrogating checkpoints and p21-mediated senescence onset (Fig. S9A). Thus, p21 appears to be the main effector mediating DNA damage-induced cell cycle exit after Chk1 depletion. In addition, our data support the role of sustained Chk1 activation in preventing p21/RB-mediated permanent G2 exit, and corroborate the importance of CDK6 downregulation in senescence onset in cancer cells.

## DISCUSSION

Our findings reveal that senescence onset upon permanent DNA damage-induced G2 arrest (G2 exit) involves interplay between the p21-cycD-RB pathway and Chk1/Chk2 kinases (Fig. 6G). In non-transformed cells G2 exit is triggered by p21-dependent RB activation that correlates with Chk1 downregulation and precedes the acquisition of senescence hallmarks such as p16 or SA-β- gal. This G2 exit programme is compromised in p53/RB-proficient cancer cells due to inefficient p21-dependent inhibition of RB phosphorylation and sustained Chk1 activation. Acute Chk1 depletion in these cells dramatically accelerates cell cycle exit, in p21-dependent manner, by inhibiting or downregulating RB kinases.

We identify CycD1-CDK2/CDK4 complexes as major p21 targets after G2 arrest, thus uncovering their previously unrecognized role in RB phosphorylation in late cell cycle phases (Chung et al, 2019; Topacio et al, 2019). Although often considered (with p27^Kip1^) as a stabiliser, and even activator, of CycD1-CDK complexes (Sherr & Roberts, 1999), p21 was shown to inhibit CycD1-CDK4 complexes (Guiley et al, 2019; Yang et al, 2017b). Therefore, we propose that p21- CycD1 is an RB-linked DNA damage effector even beyond the G1/S transition, enabling the transition from a temporary G2 arrest to a permanent G2 exit (Fig. 6G). Our results are fully consistent with the recent finding that mitogen-dependent CycD1 levels in G2 phase drive cell cycle progression (Min et al, 2020). However, we cannot exclude a possibility that p21 promotes G2 exit also by inhibiting CycA-Cdk2 that phosphorylates RB as well (Topacio et al, 2019).

In addition to its role in blocking G2/M progression, p21 targets CycD1-CDK2 and CycE1- CDK2 complexes, which accumulate upon mitotic bypass, thereby preventing re-replication (Fig. 6G) that might otherwise take place upon APC/C^Cdh1^ inactivation (Cappell et al, 2016). The importance of inhibition of CycD1-CDK2 by p21 is highlighted by the recent discovery that this complex, which efficiently phosphorylates RB (Chytil et al, 2004; Topacio et al, 2019), enables resistance of cancer cells to CDK4/6 inhibitors such as palbociclib (Chaikovsky et al, 2021; Saengboonmee & Sicinski 2021). Interestingly, in p53-proficient cancer cells, p21-bound CycD1/CycE1 complexes increase after prolonged G2 arrest, concomitant with CycA downregulation and cell cycle exit, but its significance is unclear. We speculate that these complexes, strongly upregulated after Chk1 depletion but also observed in “spontaneously” quiescent (Gookin et al, 2017) or senescent (Bacevic et al, 2017a; Stein et al, 1999) cells, might play a role in senescence, for example, by preventing apoptosis.

The notion that p21-RB-mediated Chk1 downregulation promotes permanent cell cycle exit might appear counterintuitive, since sustained Chk1 activity stabilises G2 arrest enabling senescence onset (Johmura et al, 2016). Nonetheless, Chk1 suppression has been observed after DNA damage induced cell cycle arrest or senescence (Gabai et al, 2008; Gottifredi et al, 2001) and even implicated in prevention of apoptosis during trophoblast differentiation (Ullah et al, 2011). However, given its key role in DNA damage checkpoints, Chk1 inactivation or downregulation was often regarded as a “defect in the DNA damage response” (Gabai et al, 2008), a mechanism enabling adaptation to genotoxic stress (Shaltiel et al, 2015), recovery from severe replication stress (Zhang et al, 2005), cell cycle resumption after DNA damage repair (Park et al, 2015) or preventing a prolonged G2 arrest which might trigger apoptosis (Gottifredi et al, 2001). In contrast, our results implicate Chk1 downregulation as a component of the senescence programme (Fig. 6G). Prolonged Chk1 activity might promote CDK-mediated RB inactivation and prevent senescence onset by restraining p21 induction (Beckerman et al, 2009; Hsu et al, 2019), positively regulating CDK6 expression by inhibiting E2F6, a repressor of E2F-dependent transcription (Bertoli et al, 2013) or by phosphorylating and inactivating E2F7 and E2F8, potent transcriptional repressors (Yuan et al, 2018). Moreover, the results showing that telomere dysfunction in the absence of p53 associates with sustained Chk1 activation and endoreplication (Davoli et al, 2010) are fully consistent with our model. That Chk1 activity compromises senescent onset might explain the surprising result that an extra copy promotes oncogenic transformation in mice (Lopez-Contreras et al, 2012).

Our results showing that DNA damage-induced G2 exit correlates with p53-dependent diminution of gH2AX signal and reduction of gH2AX/53BP1 foci could reflect merely an energy saving switch-off of DNA damage signalling once the cells are committed to permanent cell cycle exit. Also, this might explain low number of in gH2AX/53BP1 foci in senescent cells (Gire et al, 2004). However, it is tempting to speculate that gH2AX/53BP1 foci clustering (Aymard et al, 2017), resembling 53BP1 nuclear bodies (Lukas et al, 2011), might be involved in senescence onset.

Our finding that prolonged DNA damage or accelerated senescence after Chk1 depletion in cancer cells downregulates CDK6 but not CDK4, not only underlines differential regulation of the two kinases but also uncovers a novel, CDK inhibitor-independent mechanism of inhibiting RB phosphorylation. Thus, Chk1 might promote cell cycle progression by positively regulating CDK6 expression, for example, by inhibiting E2F6, a repressor of E2F-dependent transcription (Bertoli et al, 2013). Nonredundant role of CDK6 in RB phosphorylation is also supported by findings in *CDK6^-/-^* mice (Santamaria et al, 2007) and our recent study (Bacevic et al, 2017a). Moreover, acquired CDK6 overexpression confers cancer resistance to CDK4/6 (Yang et al, 2017a) and CDK1/2 inhibitors (Bacevic et al, 2017b). Interestingly, unlike CDK4 or CDK2, CDK6 is less targeted by p21, but upon DNA damage (this study) or in differentiation (Fujimoto et al, 2007) permanent cell cycle arrest correlates with its downregulation. Also, in replicative senescence, p16- mediated CDK6 sequestration preferentially downregulates CycD1-CDK6 but not CycD1-CDK4 complexes that are p21-bound (Dulic et al, 2000; Stein et al, 1999).

Perhaps the most surprising result is that acute Chk2 depletion did not decrease p21 induction, as expected from its role as p53 activator (Chen & Poon, 2008), but instead stimulated RB phosphorylation and Ki67 expression in G2-arrested cells. While our present and earlier results (Bacevic et al, 2017a; Lossaint et al, 2011) do not support its presumed but still debated role in G2 arrest (Stolz et al, 2011), they suggest that Chk2 might promote senescence by inhibiting RB kinases (Fig. 6G). This is consistent with its role in senescence (Chen & Poon, 2008; Gire et al, 2004) and as a global tumour suppressor associated with DNA damage (Stolz et al, 2011; Stracker et al, 2008).

In conclusion, our results support a model where reciprocal regulation of p21 and Chk1 is a core feature of a network that includes both G2/M checkpoint (Chk1, Chk2) and senescence (p21 and RB) regulators, and whose combined output controls the fate of G2-arrested cells. Therefore, inhibiting Chk1 in cancers might have therapeutic benefit in combination with genotoxic chemotherapy, as it should promote senescence of cancer cells subjected to DNA damage. As opposed to “synthetic lethality”, we propose that “synthetic senescence promotion” would be an appropriate description of such an effect.

## MATERIALS AND METHODS

### Cell lines

Normal human diploid foreskin fibroblasts (HDF), HDF expressing HPV16-E6 and human mammary epithelial cells (HMEC), from frozen stocks, were obtained, cultured and synchronized as described previously (Baus et al, 2003; Lossaint et al, 2011). U2OS (human osteosarcoma) cells were originally purchased from ATCC and HCT-116 (colon carcinoma) cells were a gift of Dr. B. Vogelstein (Johns Hopkins University, Baltimore, USA) from 2000 to 2010. The human fibroblasts (BJ) expressing inducible SV40 mutant (T_121_) specifically targeting pocket proteins (Conklin et al, 2012) were a gift from Dr. J. Sage (Stanford, USA, in 2018). Cell lines were not authenticated but were weekly tested for mycoplasma contaminations (Mycolalert kit).

Excepting for HMEC (MEGM medium; Lonza, Basel, Switzerland) and HCT-116 (McCoy’s 5A medium) all cell lines were cultured in Dulbecco modified Eagle medium (MEM (high glucose, pyruvate, GlutaMAX – Gibco® Life Technologies) supplemented with 10% foetal bovine serum (SIGMA, Dutcher or HyClone). Cells were grown under standard conditions at 37°C in humidified incubator containing 5% CO_2_.

### Cell drug treatments, radiation

Radiomimetic agent bleomycin (10 µg/ml) and topoisomerase II inhibitor ICRF-193 (bis(2,6-dioxopiperazin)), 2 µg/ml) were added to asynchronously growing cells as described previously (Baus et al, 2003). Where indicated, cells were irradiated (5 or 10 Gy) in the panoramic ^60^Co gamma irradiation facility at the Ruđer Bošković Institute (Zagreb, Croatia). Caffeine (SA, 5 mM) and ATM inhibitor KU-0055933 (Kudos Pharmaceuticals, Cambridge, UK, 10 µM) were added 1 hr before treatment with drugs. Doxycycline (ThermoFisher) was added 6-12 hr before adding the genotoxic drugs (1 μg/ml).

### Cell cycle analysis

Cell cycle analysis was determined by flow cytometry (FACS) of propidium iodide (PI)- stained cells using BD FACS Calibur (BD Biosciences, San Jose, CA) as described earlier (Bacevic et al, 2017a). Cells were harvested, washed with cold PBS, resuspended in 300 µL PBS and fixed with 700 µL ice-cold 100% methanol. Fixed cells were kept at -20°C. For the analysis, cells were pelleted by centrifugation at 6000 rpm for 5 min. After washing once with 1% BSA in PBS, cells were stained with PI staining solution (10 µg/ml PI, 1% BSA, 200 µg/ml RNase A in PBS) for 15 min at room temperature and subjected to cell cycle analysis using BD FACS Calibur (BD Biosciences, San Jose, CA). Data were analysed using FlowJo software (v10.6.1, FlowJo, LLC, Ashland, OR).

### siRNA transfection

The SMARTpool ON-TARGETplus siRNAs (CHK1, CHK2, CDK2, CDK4, CDK6 or CDKN1A-p21) were purchased from GE Dharmacon Research (Lafayette, CO, USA). As control, we used siRNA for luciferase, 5’- ACUGACGACUCUGCUACUC-3’ (Luc; Eurogentec, Seraing, Belgium) or ON-TARGETplus Non-targeting siRNA #1 (Cont; GE Dharmacon). Cells were transfected with siRNA (40 nM) using a standard calcium phosphate transfection method (Bacevic et al, 2017a). 24 hr after transfection, the cells were exposed to genotoxic agents for indicated times and harvested for biochemical or immunofluorescence analysis or monitored by video-microscopy (Lossaint et al, 2011).

Preparation of cell lysates and conditions for SDS-PAGE and immunoblotting have been described previously (Baus et al, 2003; Stein et al, 1999). Briefly, cells were harvested by trypsinisation and washed in cold PBS prior freezing in liquid N_2_. Frozen pellets (kept at -80°C) were lysed in lysis buffer (150 mM NaCl, 50 mM Tris-HCl pH 7.5, 0.2% NP-40, 2 mM EDTA, 1 mM DTT, 0.1 mM NaVO_4_, protease inhibitor cocktail (Sigma Aldrich P 8340) and incubated on ice (60 min). Lysates were centrifuged at 13,000 rpm (5 min) and supernatant was frozen (80°C). Total protein was quantified using the BCA Protein Assay Kit (Pierce Biotechnology; ThermoFisher #23227). For immunoblot analysis, cell lysates were denatured in Laemmli buffer and boiled at 95°C for 5 min. 30 μg of protein was loaded into each lane. Samples were run on 7.5%, 11% or 12.5% SDS-PAGE gels, depending on the proteins of interest, and transferred to Immobilon membranes (Merck Millipore, Burlington, MA, USA) using Owl HEP-1 semidry electroblotter (ThermoFisher). Membranes were routinely stained with Naphtol blue black (Amido-black; Sigma-Aldrich 3393) to verify the transfer and loading. Primary antibodies were diluted in 5% milk in TBS-T and incubated for 2 hr at RT or overnight at 4°C. Secondary antibodies (goat anti-mouse IgG-HRP, DACO, Glostrup, Denmark and donkey anti-rabbit IgG-HRP, GE Healthcare) were diluted 1:5000 in 5% milk in TBS-T and incubated for 1 hr at room temperature. Chemiluminescence was detected using Western Lightning Plus/Ultra (PerkinElmer, France) and Amersham Hyperfilm^TM^ (GE Healthcare).

### Antibodies

Primary antibodies for western blotting: cyclin D1 (Santa Cruz Biotechnology (SCBT) DCS-6, sc-20044), cyclin D2 (SCBT, sc-593), cyclin D3 (SCBT, sc-182), cyclin E1 (SCBT sc-247), cyclin A (6E6, Novocastra), cyclin B1 (SCBT, sc-752; 1:100), CDK1 (BD Transduction Laboratories (BDTL) C12720), CDK2 (Abcam ab128167), CDK4 (SCBT sc-260; 1:1000), CDK6 (SCBT sc-177), p16 (BD Pharmingen 550834; 1:100), p21 (SCBT sc-397 and Cell Signaling Technology (CST) 2946, 2947), p27 (SCBT sc-528; and BDTL K25020), RB (BD Pharmingen 554136; 1:200), p130 (SCBT sc-317; 1:200), RB phospho-S780 (CST 9307), Chk1 (SCBT sc-8408), Chk2 (SCBT sc-17747), Chk1 phospho-S317 (CST 12302), Chk1 phospho-S345 (CST 2348), Chk2 phospho-T68 (CST 2661), Cdc25C phospho-S216 (CST 4901), p53 (SCBT sc-126), p53 phospho-S15 (CST 9284), Ki-67 (Abcam ab16667). Dilutions were 1:500 or 1:200 for phospho-proteins, unless otherwise specified. For more details see Supplementary information for antibodies.

### Co-immunoprecipitation, p21 immuno-depletion

Cells were lysed as described above. Routinely, 100-200 μg of total cell protein was used per IP and antibody against target protein was added at a concentration of 4 μg antibody/mg of total protein, unless otherwise specified. Samples were incubated with the primary antibody for 2 hr at 4°C and the immuno-complexes were recovered using Protein A or Protein G Sepharose beads (Amersham Biosciences - 10 μl/100 ml of lysates). Beads were washed in TBS-T, and immunoprecipitated proteins were eluted in 2x Laemmli buffer by heating at 37°C for 15 min. To detect proteins on immunoblots containing immunoprecipitates peroxidase-conjugated protein A/G was used (Pierce Biotechnology, Rockford, IL, USA 32490). Primary antibodies for immunoprecipitations: cyclin E1 (SCBT sc-248), p21 (SCBT sc-397), p27 (SCBT sc-528) and rabbit polyclonal anti-cyclin D1 (Dulic et al, 1993) 1 μl/100 ml of lysate). For more details see Supplementary information for antibodies.

For p21 or p27 immunodepletion experiments, cell lysates (100-200 μg) were incubated with saturating amounts of p21 or p27 antibodies, whereas mock samples were incubated with protein A-Sepharose only. The resulting supernatants were analysed by immunoblotting as described previously (Stein et al, 1999).

### Immunofluorescence, image analysis

Experimental conditions for immunofluorescence, primary antibodies and image analysis (image acquisition and analysis) have been published previously (Bacevic et al, 2017a; Lossaint et al, 2011). Briefly, cells were seeded on coverslips and fixed in cold 100% methanol (10 min, - 20°C) or formaldehyde (3.7%, 15 min, RT). Prior to incubation with primary antibodies (1-2 hr at RT in humidified chamber) formaldehyde-fixed cells were permeabilized in 0.2% Triton X-100. Primary and secondary antibodies were diluted in blocking solution (0.1% Tween-20 in PBS with 5% FCS, 2 hr/RT). After washing in PBS-Tween (3 times for 5 min) the cells were incubated with secondary antibody (1 hr at RT) and washed again (3 times for 5 min). Secondary antibodies were diluted 1:1000 for fluorophores AlexaFluor 488, 555, and 568, and 1:500 for AlexaFluor 647 (Invitrogen, Fisher Scientific). Coverslips were rinsed in distilled water prior to mounting on slide with ProLong Diamond Antifade Mountant with DAPI (Molecular Probes P36962).

Immunofluorescence images were captured on microscope Leica CTR6000 (objective Leica 40X HCX PL APO 1.25-0.75 oil, camera CollSnap HQ2) driven by MetaMorph (MDS, Analytical Technologies, Canada). Composites were generated using Adobe Photoshop (Adobe systems, Inc, San Jose, CA, USA) and Microsoft PowerPoint (Microsoft Corp, Redmond, WA, USA) softwares. For the panels showing immunofluorescence images, a representative field was shown.

Following primary antibodies were used: cyclin E1 (HE12, SCBT sc-247; 1:100), cyclin D1 (CST 2926; Abcam ab16663), cyclin A (Novocastra 6E6 and SCBT sc751), cyclin B1 (SCBT, sc-752; 1:100), p21 (CST 2946, 2947), H2A.X phospho-S139 (MerckMillipore clone JBW301, 05- 636), 53BP1 (Novus biologicals NB100-304); Ki-67 (Abcam ab16667 and BDTL 610968).

Dilutions were 1:500 unless otherwise specified. For more details see Supplementary information for antibodies.

Every experiment was performed at least twice with each genotoxic agent. For each condition, at least ten images (40X magnification) were taken with a widefield fluorescent microscope for each situation. Immunofluorescence signals were quantified using a home-made macro in ImageJ software (Virginie Georget, MRI) as described previously (Bacevic et al, 2017a). DAPI images were used to identify the individual nuclei using a background correction, a mask creation based on threshold, watershed segmentation and the “analyse particles” function in ImageJ. The region-of-interest corresponding to the nuclei was applied to the different channels and total intensity of individual nuclei was quantified. The boxplots from quantification data were generated with R (version 3.6.2) and represent cell pools of at least two independent experiments. For the panels showing immunofluorescence images, representative fields were shown.

### Video-microscopy

Conditions for video-microscopy were described previously (Bacevic et al, 2017a; Lossaint et al, 2011). Mitoses were scored by inspection of video-microscopy sequences (MetaMorph software). Images were taken at 10-15 min intervals for at least 48 hours. Three fields for each situation were analysed and normalized for the cell number at the beginning of the time-lapse sequence. For mitosis-entry kinetics, total number of mitotic cells during the given interval was plotted. For each experiment, all the conditions were tested in parallel, including controls with untreated cells transfected with different siRNAs.

### SA-β-galactosidase

Conditions for SA-β-galactosidase staining using Senescence detection kit (Abcam, ab655351) were described previously (Bacevic et al, 2017a; Sobecki et al, 2017). HDF or U2OS cells were seeded on coverslips in 12-well plate, at density of 10^5^ cells per well. After 24 hr, cells were treated with ICRF-193 (2 µg/ml) or bleomycin (10 µg/ml). The drug was washed away from the cells after 48 hr incubation, followed by staining after 2 or 4 weeks using Senescence detection kit (Abcam, ab655351). Photos were taken using an upright microscope at 20X magnification.

### Statistical analysis

Each experiment was performed independently two to three times. Data are presented as the mean ± standard deviation (SD). The two-sided Student’s *t*-test was performed using Microsoft^®^ Excel^®^ 2016 (Microsoft Corp, Redmond, WA, USA) to analyse the differences between the means of groups. Differences were considered statistically significant for a *p* value of ≤0.05 labelled by an asterisk (*). Immunofluorescence images were quantified using ImageJ software. Box plots and violin plots were generated using R software. For all analyses, the cells were pooled from two to three independent experiments and at least 200 cells were analysed.

## SUPPORTIG INFORMATION

Supporting information includes nine figures and one list of antibodies used.

## ACKNOWLEDGEMENTS

This work was supported by grants of ARC (N°3793 to V.D.) and the Cancéropole du Grand Sud Ouest. The team (GL, KB, VD, KM and DF) is “Equipe labellisé” by the Ligue Nationale Contre le Cancer (EL2013.LNCC/DF). GL and KB were recipients of a PhD fellowship from LNCC. A.H. was funded by Croatian Science Foundation (grant IP-2013-11-1615). V. Gire was funded by INCA 2017-169. We thank Dr A. Péléraux for statistical analysis of immunofluorescence images and the figures using R software. We thank Drs E. Schwob, A. Camasses, J. Piette, J. Sage and P. Coulombe for critically reading the manuscript. We are grateful to KUDOS Pharmaceuticals for gift of KU-55933 and Dr. J. Sage (Stanford, USA) for generous gift of T_121_-expressing fibroblasts. We acknowledge the imaging facility MRI, member of the national infrastructure France-BioImaging supported by the French National Research Agency (ANR-10-INBS-04, «Investments for the future»).

## AUTHOR CONTRIBUTIONS

VD conceived and supervised the project, VD, GL and AH designed research and interpreted the data, GL, AH, VGi, KB, VD and KM performed the experiments. VGe created ImageJ macros for immunofluorescence image quantifications. DF and VD wrote the manuscript. All authors reviewed the manuscript.

## Supplementary information for antibodies used

### Antibodies for western blotting

cyclin D1 (Santa Cruz Biotechnology (SCBT) DCS-6, sc-20044), https://www.scbt.com/fr/p/cyclin-d1-antibody-dcs-6

cyclin D2 (SCBT, sc-593), https://www.scbt.com/fr/p/cyclin-d2-antibody-m-20

cyclin D3 (SCBT, sc-182) https://www.scbt.com/p/cyclin-d3-antibody-c-16

cyclin E1 (HE12, SCBT sc-247), https://www.scbt.com/fr/p/cyclin-e-antibody-he12

cyclin A (6E6, Novocastra), https://www.citeab.com/antibodies/725566-ab16726-anti-cyclin-a2-antibody-6e6

cyclin B1 (GNS1, SCBT, sc-752; 1:100), https://www.scbt.com/p/cyclin-b1-antibody-gns1?gclid=EAIaIQobChMIibjigZnR8QIVygyLCh213wWrEAAYAiAAEgLh9fD_BwE

<u>CDK1 (BD Transduction Laboratories (BDTL) C12720)</u>, https://www.bdbiosciences.com/en-us/products/reagents/microscopy-imaging-reagents/immunofluorescence-reagents/purified-mouse-anti-cdk1.610037

CDK2 (Abcam ab128167), https://www.abcam.com/cdk2-antibody-1a6-ab128167.html

CDK4 (SCBT sc-260; 1:1000), https://www.scbt.com/fr/p/cdk4-antibody-c-22

CDK6 (SCBT sc-177), https://www.scbt.com/fr/p/cdk6-antibody-c-21

p16 (BD Pharmingen 550834), https://www.bdbiosciences.com/ds/pm/tds/550834.pdf

p21 (SCBT sc-397 and Cell Signaling Technology (CST) 2946, 2947), https://www.scbt.com/fr/p/p21-antibody-c-19

https://www.cellsignal.com/products/primary-antibodies/p21-waf1-cip1-12d1-rabbit-mab/2947

https://www.cellsignal.com/products/primary-antibodies/p21-waf1-cip1-dcs60-mouse-mab/2946

p27 (SCBT sc-528; and BD Transduction Laboratories (BDTL K25020), https://www.scbt.com/fr/p/p27-antibody-c-19

https://www.bdbiosciences.com/en-us/products/reagents/microscopy-imaging-reagents/immunofluorescence-reagents/purified-mouse-anti-p27-kip1.610242

<u>RB (BD Pharmingen 554136),</u> https://www.bdbiosciences.com/en-us/products/reagents/flow-cytometry-reagents/research-reagents/single-color-antibodies-ruo/purified-mouse-anti-human-retinoblastoma-protein.554136

p130 (SCBT sc-317), https://www.scbt.com/fr/p/p130-antibody-c-20

RB phospho-S780 (CST 9307), https://www.cellsignal.com/products/primary-antibodies/phospho-rb-ser780-antibody/9307

Chk1 (SCBT sc-8408), https://www.scbt.com/fr/p/chk1-antibody-g-4

Chk2 (SCBT sc-17747), https://www.scbt.com/fr/p/chk2-antibody-a-11

Chk1 phospho-S317 (CST 12302), https://www.cellsignal.com/products/primary-antibodies/phospho-chk1-ser317-antibody/2344

Chk1 phospho-S345 (CST 2348), https://www.cellsignal.com/products/primary-antibodies/phospho-chk1-ser345-133d3-rabbit-mab/2348

Chk2 phospho-T68 (CST 2661), https://www.cellsignal.com/products/primary-antibodies/phospho-chk2-thr68-antibody/2661

Cdc25C phospho-S216 (CST 4901), https://www.cellsignal.com/products/primary-antibodies/phospho-cdc25c-ser216-antibody/9528

p53 (SCBT sc-126), https://www.scbt.com/p/p53-antibody-do-1?gclid=EAIaIQobChMIrLCO1Z7R8QIVq4ODBx0oNwKBEAAYASAAEgJQkvD_BwE

<u>p53 phospho-S15 (CST 9284)</u>, https://www.cellsignal.com/products/primary-antibodies/phospho-p53-ser15-antibody/9284

Ki-67 (Abcam ab16667) https://www.abcam.com/ki67-antibody-sp6-ab16667.html

### Co-immunoprecipitation

Primary antibodies for immunoprecipitations: cyclin E1 (HE111, SCBT sc-248),

https://www.scbt.com/p/cyclin-e-antibody-he111

p21 (SCBT sc-397),

https://www.scbt.com/fr/p/p21-antibody-c-19

p27 (SCBT sc-528),

https://www.scbt.com/fr/p/p27-antibody-c-19

rabbit polyclonal anti-cyclin D1

https://www.ncbi.nlm.nih.gov/pmc/articles/PMC47916/

### Immunofluorescence

cyclin E1 (HE12, SCBT sc-247),

https://www.scbt.com/fr/p/cyclin-e-antibody-he12

cyclin D1 (CST 2926, Abcam ab16663),

https://www.cellsignal.com/products/primary-antibodies/cyclin-d1-dcs6-mouse-mab/2926

https://www.abcam.com/cyclin-d1-antibody-sp4-ab16663.html

cyclin A (Novocastra 6E6 and SCBT sc751),

https://www.citeab.com/antibodies/725566-ab16726-anti-cyclin-a2-antibody-6e6 https://www.scbt.com/fr/p/cyclin-a-antibody-h-432

p21 (CST 2946, 2947),

https://www.cellsignal.com/products/primary-antibodies/p21-waf1-cip1-12d1-rabbit-mab/2947

https://www.cellsignal.com/products/primary-antibodies/p21-waf1-cip1-dcs60-mouse-mab/2946

H2A.X phospho-S139 (Merck Millipore clone JBW301, 05-636),

https://www.merckmillipore.com/FR/fr/product/Anti-phospho-Histone-H2A.X-Ser139-Antibody-clone-JBW301,MM_NF-05-636-I?ReferrerURL=https%3A%2F%2Fwww.google.com%2F&bd=1

53BP1 (Novus biologicals NB100-304);

https://www.novusbio.com/products/53bp1-antibody_nb100-304

Ki-67 (Abcam ab16667)

https://www.abcam.com/ki67-antibody-sp6-ab16667.html

Ki-67 (BDTL 610968)

https://www.bdbiosciences.com/en-us/products/reagents/microscopy-imaging-reagents/immunofluorescence-reagents/purified-mouse-anti-human-ki-67.610969

---

**Fig. S1.**
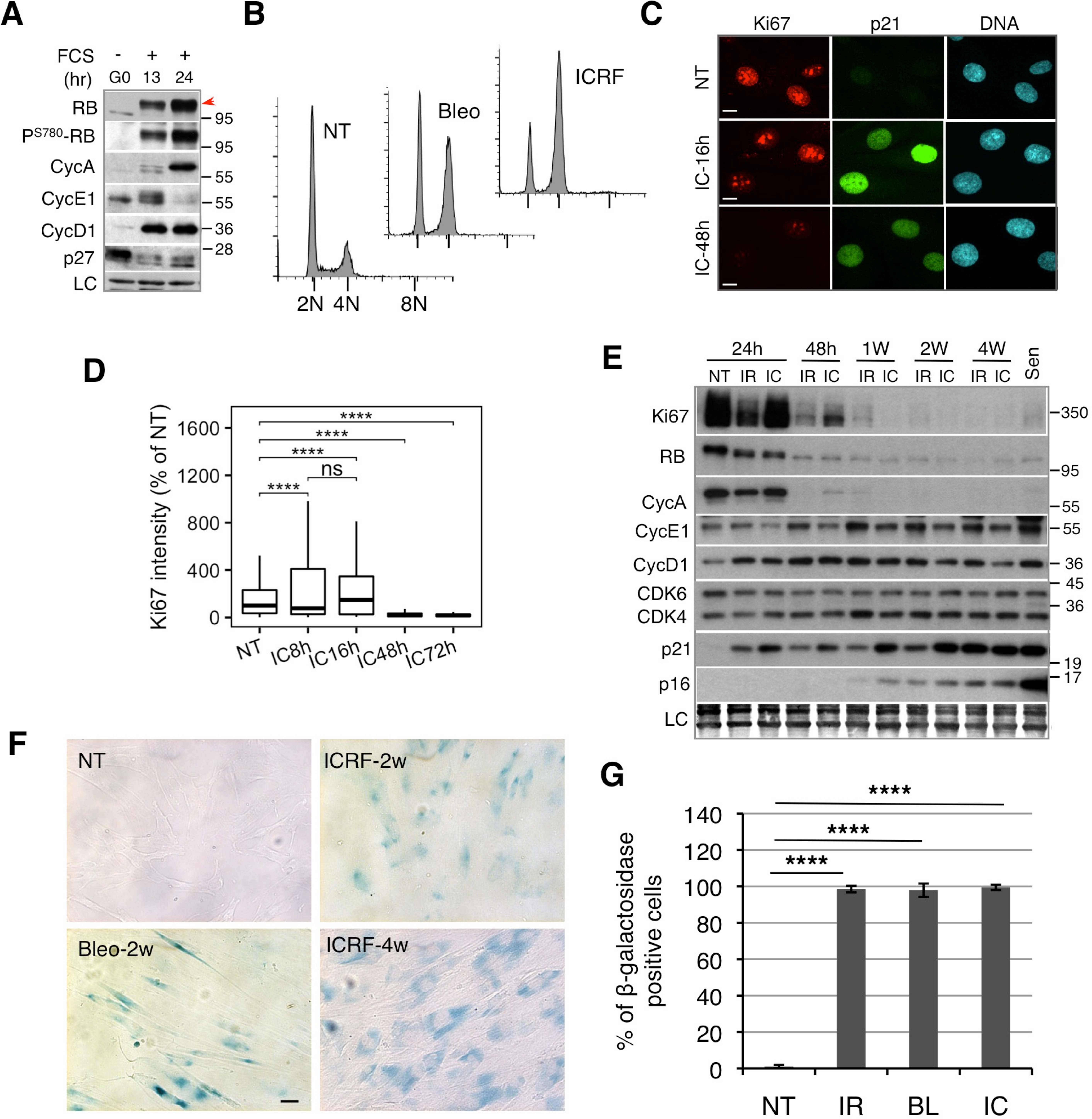
DNA damage-induced cell cycle arrest in G2 leads to cell cycle exit and senescence. **A.** Immunoblots showing RB phosphorylation (red arrow and PS780-RB) and expression of different cyclins in HDF that were released from quiescence (G0) by serum addition (+FCS) for indicated times. LC, loading control. **B.** Flow cytometry DNA content profiles of the asynchronously growing non-treated (NT) and HDF exposed to bleomycin (Bleo) or ICRF-193 (ICRF) for 48 hours. **C.** Representative immunofluorescence images (n=3) showing co-expression of Ki67 and p21 in non-treated (NT) and HDF exposed to ICRF-193 (IC) for 16 and 48 hr. Scale bar, 10 μM. **D.** Quantification of Ki67 intensity in immunofluorescence images from HDF exposed to ICRF-193 (IC) for indicated times (% of NT). Pooled cells from four independent experiments. More than 100 cells were analyzed in each experiment. NT, non-treated cells. Box plot whiskers indicate 10-90% boundary. *P* values were calculated with two-sided Student’s *t* test; ***P ≤ 0.001, ****P ≤ 0.0001. **E.** Immunoblots showing changes in cell cycle regulators in HDF upon g irradiation (IR, 10Gy) or treatment with ICRF-193 (IC) for indicated times. NT, non-treated cells; SEN, senescent cells. LC, loading control. **F.** Phase-contrast images showing β-galactosidase staining of non-treated (NT) and HDF exposed to ICRF-193 (ICRF) for 2 or 4 weeks (n=2). NT, non-treated cells. **G.** Quantification of β-galactosidase staining in HDF exposed to γ irradiation (IR, 10Gy), ICRF-193 (IC) or bleomycin (Bl) for two weeks. NT, non-treated cells. Data are mean +/- SEM of at least two independent experiments. *P* values were calculated with two-sided Student’s *t* test; ****P ≤ 0.0001. Loading controls (LC) were amido-black stained membranes.

**Fig. S2.**
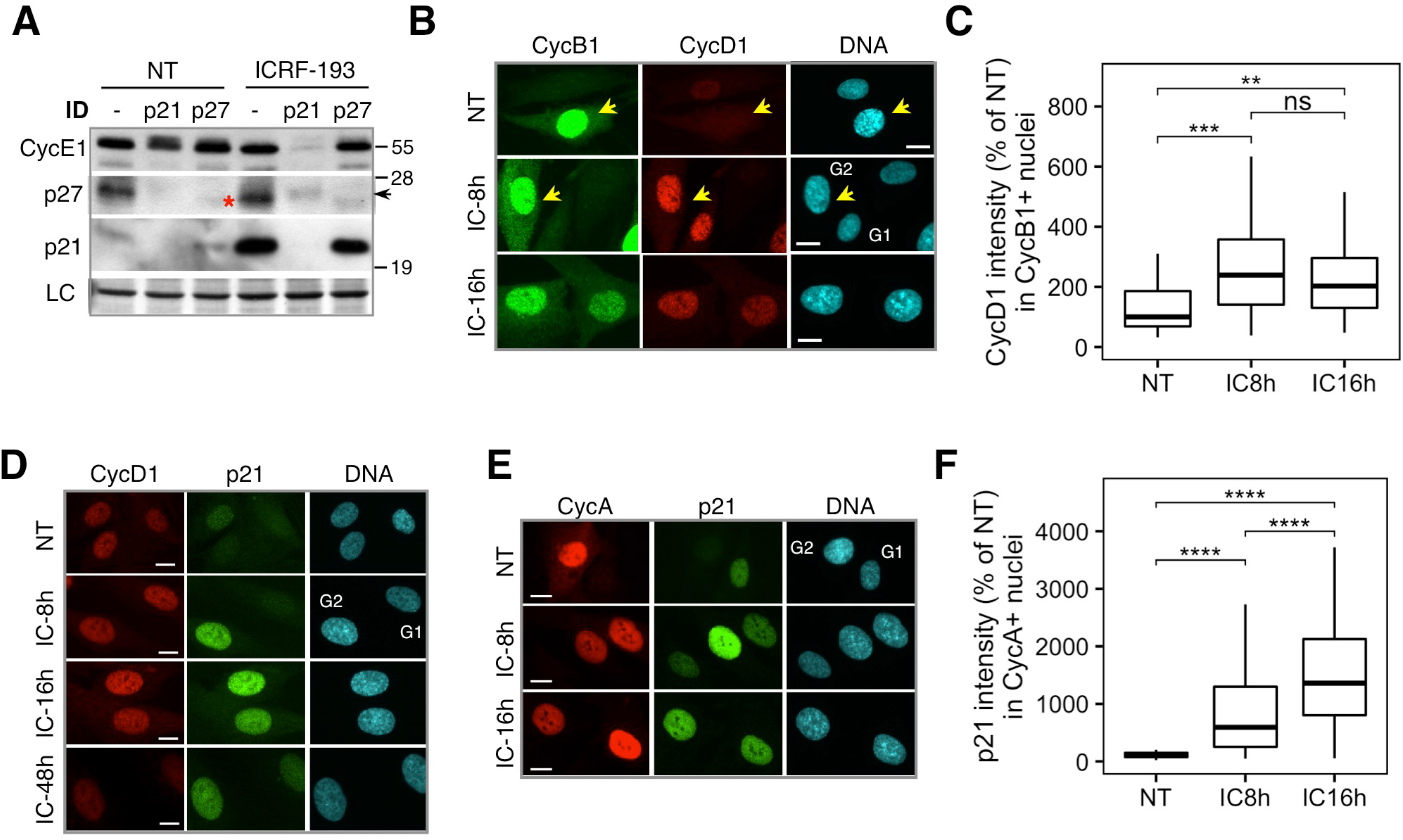
After G2 arrest p21 is co-expressed with Cyclin A and Cyclin D. **A.** p21 immunodepletion (ID) experiment. Immunoblots showing CycE1 levels before (mock-treated: -) or after immunodepletion (ID) of p21 or p27 from the lysates prepared from NT (non-treated) and HDF exposed to ICRF-193 (24h). LC, loading control. Arrow indicates position of p27 band. *, immunoblot artefact. **B.** Representative immunofluorescence images (n=2) showing co-localization of CycB1 and CycD1 in control non-treated (NT) and HDF exposed to ICRF-193 (IC) for 8h and 16h. Bar, 10 μM. Arrows points at cells expressing nuclear CycB1. **C.** Quantification of CycD1 intensity in CycB1-positive HDF exposed to ICRF-193 (IC) for indicated times. Pooled cells (n>100) from two independent experiments. Box plot whiskers indicate 10-90% boundary. **D.** Representative immunofluorescence images (n=2) showing co-localization of p21 and CycD1 in HDF exposed to ICRF-193 (IC) for indicated times. Bar, 10 μM. Quantification in Figure 1E. **E.** Representative immunofluorescence images (n=2) showing co-localization of CycA and p21 in HDF exposed to ICRF-193 (IC) for indicated times. Bar, 10 μM. **F.** Quantification of p21 intensity in CycA1-positive nuclei in HDF exposed to ICRF-193 for indicated times. Pooled cells (n>100) from two independent experiments. Box plot whiskers indicate 10-90% boundary. *P* values were calculated with two-sided Student’s *t* test; **P ≤ 0.01, ***P ≤ 0.001, ****P ≤ 0.0001.

**Fig. S3.**
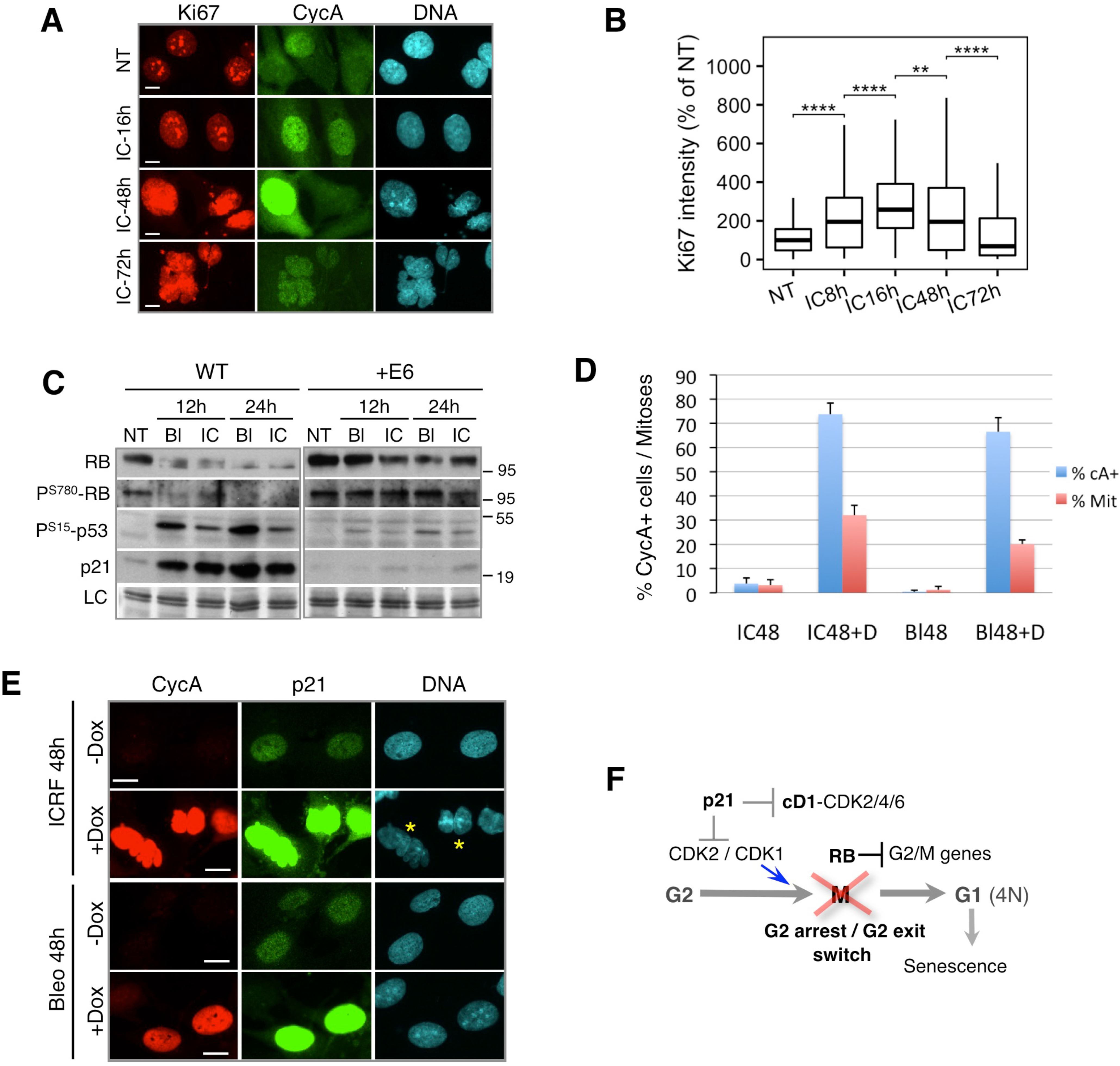
HPV-E6-mediated p53 degradation and inactivation of pocket proteins by the T_121_ mutant of SV40 oncogene compromises G2 exit in human fibroblasts. **A.** Representative immunofluorescence images (n=2) showing expression of Ki67 and CycA in HDF-E6 exposed to ICRF-193 (IC) for indicated times. Bar, 10 μM. **B.** Quantification of Ki67 intensity in the nuclei of HDF-E6 exposed to ICRF-193) for indicated times. Pooled cells (n>100) from two independent experiments. Box plot whiskers indicate 10-90% boundary. **P ≤ 0.01, ****P ≤ 0.0001. NT, non-treated cells. **C.** Immunoblots showing CycD1-specific RB phosphorylation (PS780), p53 phosphorylation (PS15) and p21 induction in WT and HPV16-E6 expressing HDF exposed to bleomycin (Bl) or ICRF-193 (IC) for 12 and 24 hours. LC, loading control. **D.** CycA positive (cA+) cells and mitotic cells (Mit, DAPI) were scored from immunofluorescence images taken at the indicated time points after the drug addition. T_121_- expressing fibroblasts were exposed to bleomycin (Bl) or ICRF-193 (IC) for 48 hr. Doxycycline (D) was added 12 hr prior to drug treatment (+). More than 100 cells were analyzed in each experiment. Data are mean +/- SEM of at least two independent experiments. **E.** Representative immunofluorescence images (n=2) showing co-expression of CycA and p21 in the presence of ICRF-193 (ICRF) and bleomycin (Bleo) in T_121_-expressing fibroblasts. Asterisks denote binuclear cells generated by unsuccessful cytokinesis after mitosis. Experimental conditions were as above. Bar, 10 μM. **F.** Model proposing the role of p21-mediated CycD1-CDK2/4 inhibition in the G2 arrest / G2 exit switch preceding senescence. Loading controls (LC) were amido-black stained membranes. *P* values were calculated with two-sided Student’s *t* test; ***P ≤ 0.001, ****P ≤ 0.0001.

**Fig. S4.**
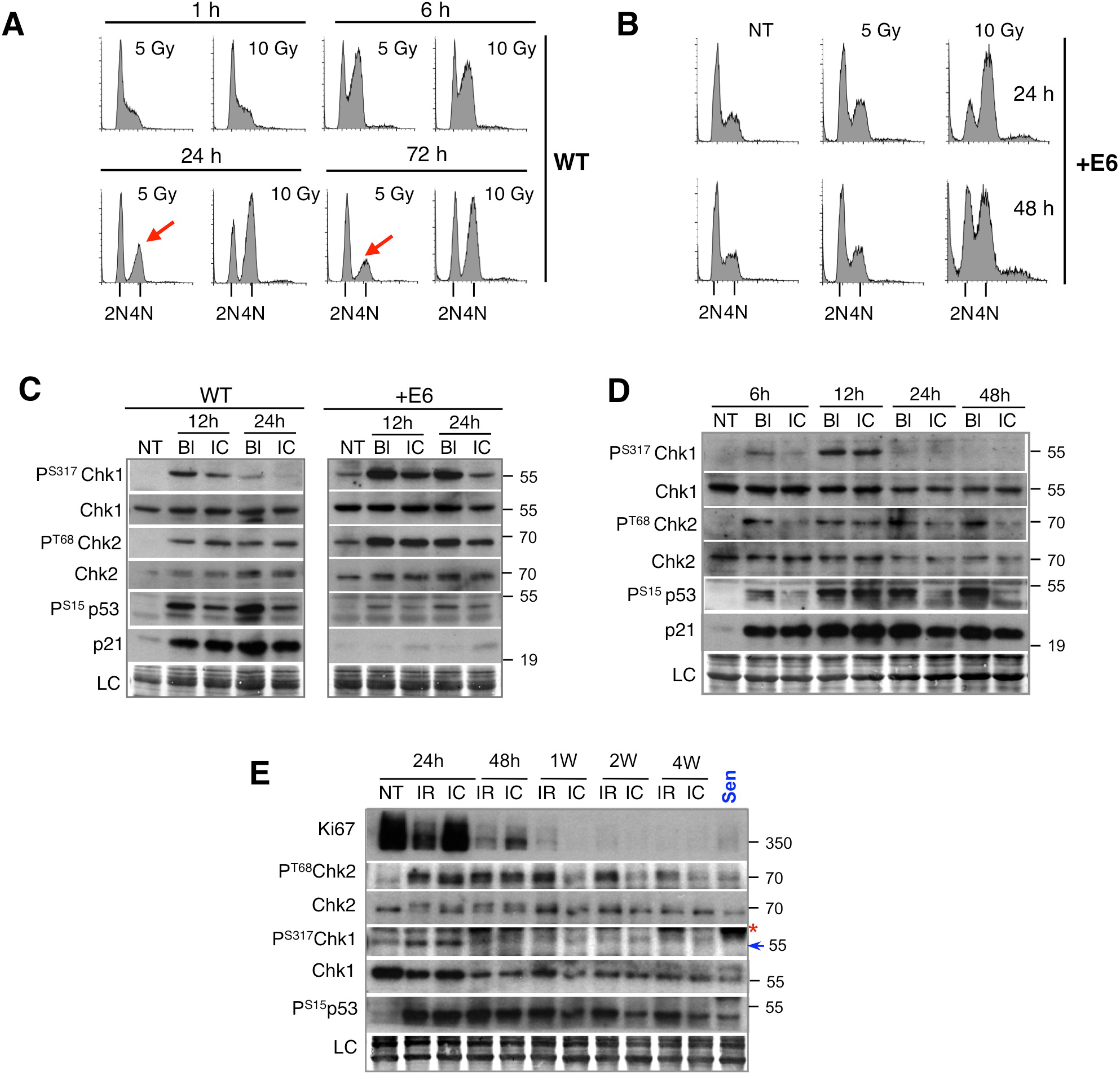
G2 exit correlates with p53-dependent suppression of Chk1 signaling. **A.** Flow cytometry DNA content of wild-type (WT) HDF at indicated times after irradiation. Cells were irradiated (5 or 10 Gy) 16 hr after release from quiescence (contact inhibition) by replating. **B.** Flow cytometry DNA of HDF expressing HPV16-E6 (+E6) 24 and 48 hours after irradiation. Asynchronous cells were exposed to 5 or 10 Gy. NT, non-treated cells. **C.** Immunoblots showing phosphorylation of Chk1 (PS317), Chk2 (PT68) and p53 (PS15) as well as p21 induction in wild-type (WT) and HDF expressing HPV16-E6 (+E6) exposed to ICRF193 (IC) or bleomycin (Bl) for indicated times. NT, non-treated cells; LC, loading control. **D.** Immunoblots showing down-regulation of Chk1 phosphorylation (P^S317^-Chk1) and sustained Chk2 (P^T68^-Chk2) phosphorylation in HDF exposed to ICRF-193 (IC) or bleomycin (Bl) at indicated times. NT, non-treated cells; LC, loading control. **E.** Immunoblots showing Ki67 expression and denoted DNA damage signalling effectors in HDF upon g irradiation (IR, 10 Gy) or treatment with ICRF-193 (IC) for indicated times. NT, non-treated cells; Sen, senescent cells. LC, loading control. Arrow: P^S317^-Chk1 band; Red asterisk: nonspecific band. Loading controls (LC) were amido-black stained membranes.

**Fig. S5.**
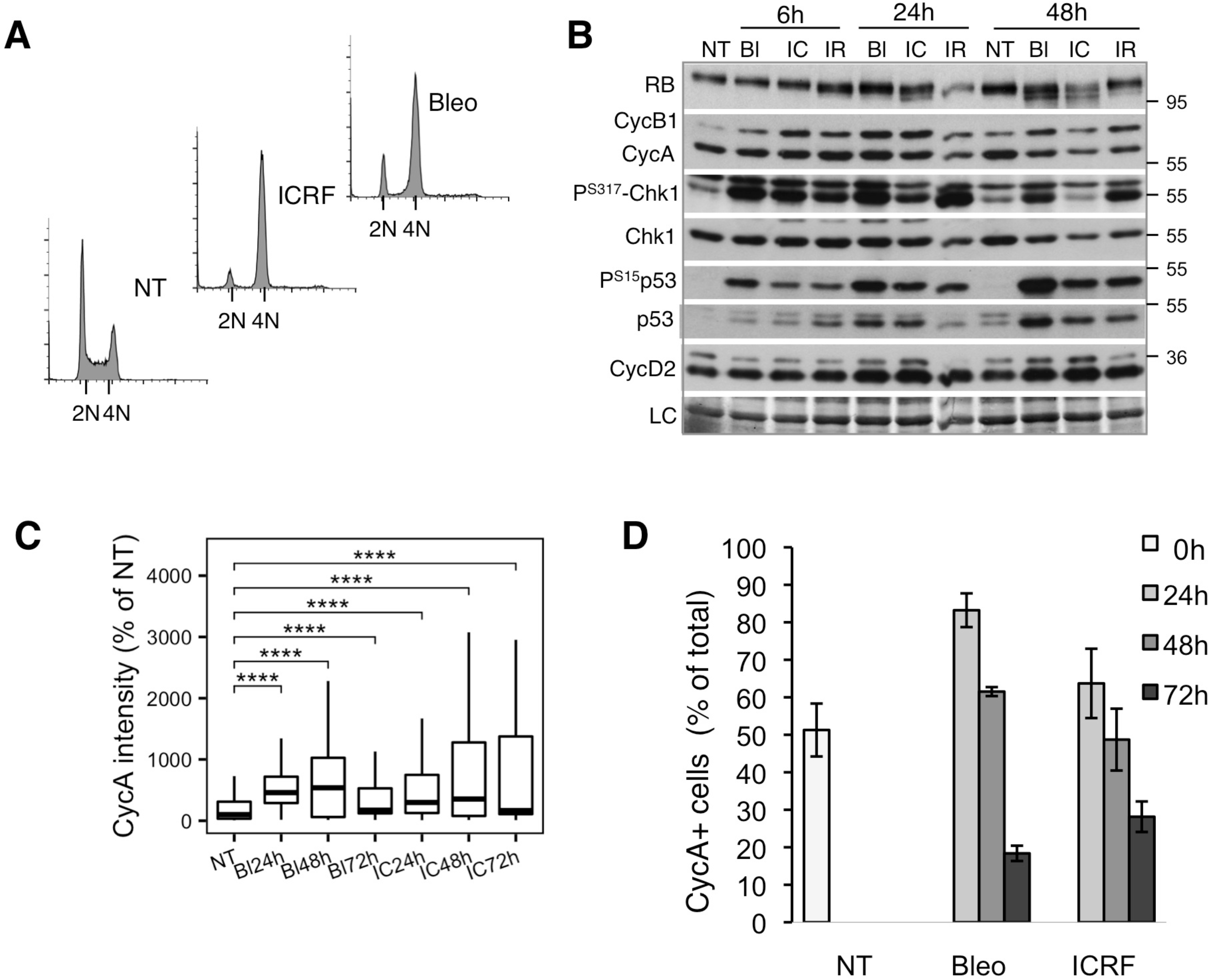
DNA damage-induced G2 arrest in U2OS cells associates with impaired cell cycle exit. **A.** Flow cytometry DNA content profiles of non-treated (NT) and U2OS cells exposed to bleomycin (Bleo) or ICRF-193 (ICRF) for 24 hr. **B.** Immunoblot analysis showing DNA damage response (P^S317^Chk1, P^S15^p53), RB phosphorylation and expression of different cyclins in U2OS cells after γ-radiation (IR, 10 Gy) or exposure to bleomycin (Bl) or ICRF-193 (IC) for indicated times. NT, non-treated cells. LC, loading control. **C.** Quantification of nuclear CycA intensity in U2OS cells exposed to bleomycin (BL) or ICRF-193 (IC) for indicated times. Pooled cells (n>100) from two independent experiments. Box plot whiskers indicate 10-90% boundary. NT, non-treated cells. *P* values were calculated with two-sided Student’s *t* test; ****P ≤ 0.0001. **D.** Quantification of CycA positive cells (% of total cell numbers) in immunofluorescence images taken at the indicated time-points after exposure to bleomycin (Bleo) or ICRF-193 (ICRF). Data are mean +/- SEM of three independent experiments. At least 100 cells were analyzed for each experiment. Loading controls (LC) were amido-black stained membranes.

**Fig. S6.**
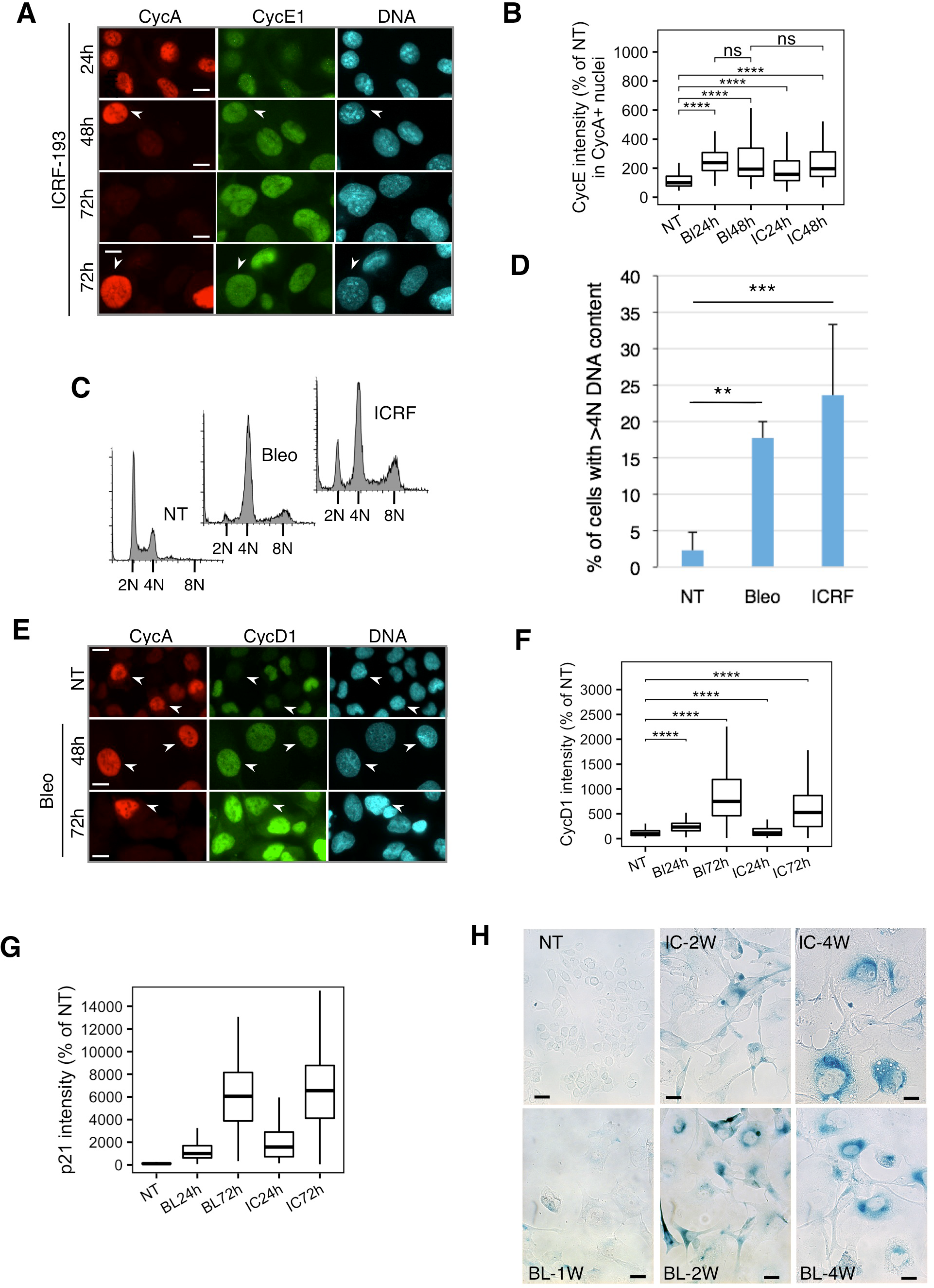
DNA damage-induced G2 arrest in U2OS leads to aberrant upregulation of G1 cyclins and p21, genome re-duplication and delayed senescence onset. **A.** Representative immunofluorescence images (n=2) showing CycA/CycE1 co-expression in U2OS cells exposed to ICRF-193 for indicated times. Arrowheads show CycA-positive cells with elevated CycE1 expression. Bar, 10 μM. **B.** Quantification of CycE1 intensity in CycA-positive nuclei in non-treated (NT) or U2OS cells exposed to bleomycin (BL) or ICRF-193 (IC) for 24 or 48 hours. Pooled cells (n>200) from two independent experiments. Box plot whiskers indicate 10-90% boundary. ****P ≤ 0.0001 **C.** Representative flow cytometry analysis of DNA content of non-treated (NT) and U2OS cells exposed to bleomycin and ICRF-193 (ICRF) for 48 hr. **D.** Percent of cells with >4N DNA content in non-treated (NT) and U2OS cells exposed to bleomycin and ICRF-193 (ICRF) for 48 hrs (n=5). Data are mean +/- SEM of five independent experiments. *P* values were calculated with two-sided Student’s *t* test; **P ≤ 0.01, ***P ≤ 0.001. **E.** Representative immunofluorescence images (n=2) showing CycA/CycD1 co-expression in non-treated (NT) and U2OS cells exposed to bleomycin (Bleo) for indicated times. Arrowheads show CycA-positive cells with elevated CycD1 levels. Bar, 10 μM. **F.** Quantification of nuclear CycD1 intensity in U2OS cells exposed to ICRF-193 or bleomycin for indicated times. Pooled cells (n>200) from two independent experiments. Box plot whiskers indicate 10-90% boundary. ****P ≤ 0.0001. NT, non-treated cells. **G.** Quantification of nuclear p21 intensity in U2OS cells exposed to bleomycin (BL) or ICRF- 193 (IC) for indicated times. Pooled cells (n>200) from two independent experiments. Box plot whiskers indicate 10-90% boundary. ****P ≤ 0.0001. NT, non-treated cells. **H.** Quantification of p21 intensity in CycE1-positive nuclei in U2OS cells not treated (NT) or exposed to bleomycin (Bl) or ICRF-193 (IC) for 24 or 48 hours. Pooled cells (n>200) from two independent experiments. Box plot whiskers indicate 10-90% boundary. **I.** Representative phase-contrast images showing β-galactosidase staining of non-treated (NT) and U2OS cells exposed to ICRF-193 (ICRF) or bleomycin (Bleo) for 1, 2 or 4 weeks (W). Bar: 10 μM.

**Fig. S7.**
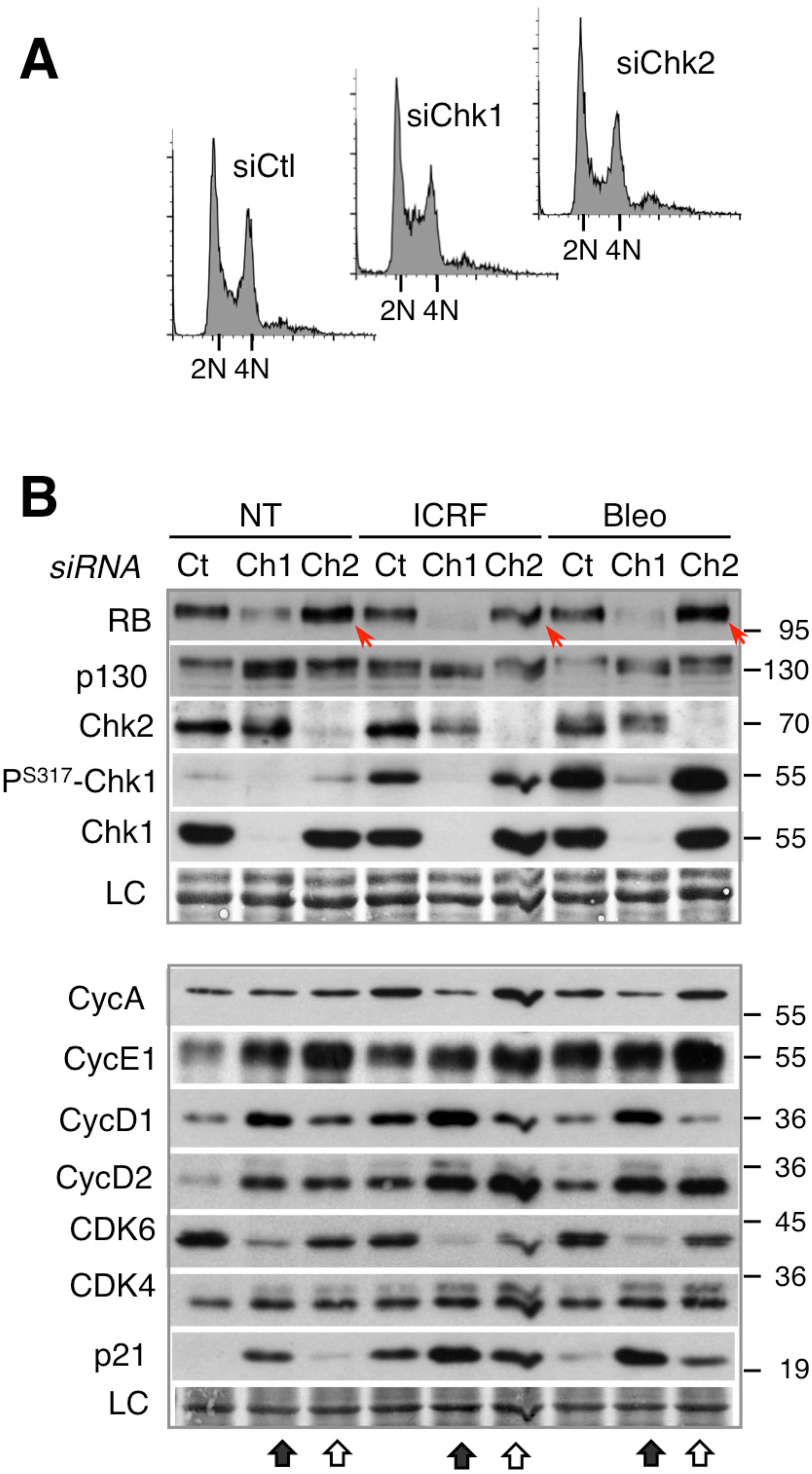
Acute Chk1 depletion promotes whereas Chk2 depletion inhibits DNA damage-induced cell cycle exit. **A.** Flow cytometry DNA content profiles of asynchronously growing control (siCtl) and U2OS cells 24 hr after siRNA-mediated Chk1 (siChk1) or Chk2 (siChk2) knockdown. **B.** Immunoblots showing effects of Chk1 (Ch1, black arrow) and Chk2 (Ch2, white arrow) knockdown on RB and p130 phosphorylation (upper panel) and expression of cell cycle regulators (lower panel) in non-treated (NT) and U2OS cells exposed to ICRF-193 (ICRF) or bleomycin (Bleo) for 16 hr. Red arrows indicate increased levels of phosphorylated RB after Chk2 knockdown. Loading controls (LC) are amido-black stained membranes.

**Fig. S8.**
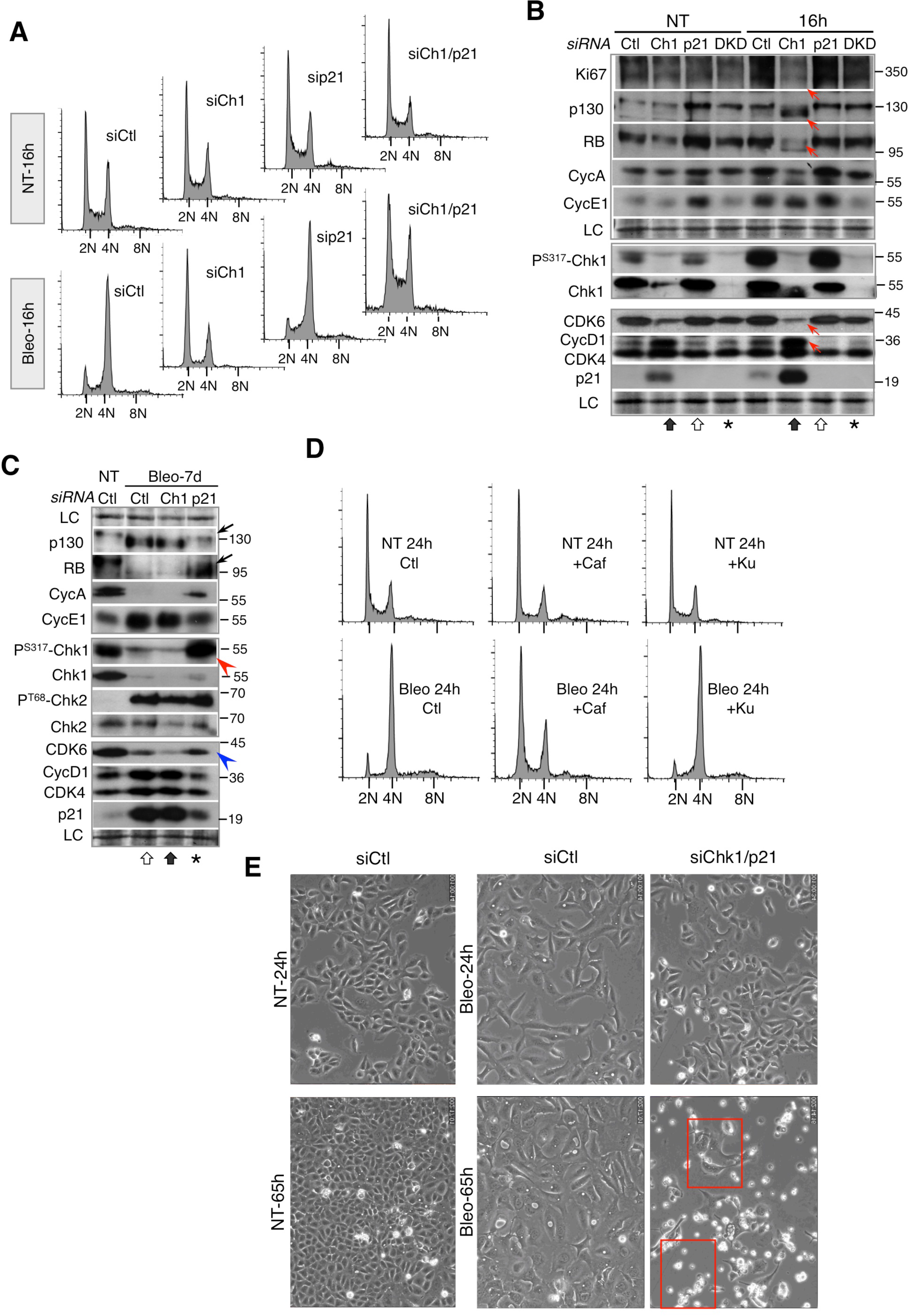
Accelerated DNA damage-induced cell cycle exit by Chk1 knockdown in U2OS cells is mediated by p21. **A.** Flow cytometry DNA content profiles of non-treated control (siCtl) and bleomycin-treated U2OS cells depleted for Chk1 (siCh1), p21 (sip21) or both proteins (siCh1/21) after 16 hr. **B.** Immunoblots showing effects of siRNA-mediated Chk1 (Ch1), p21 or double p21/Chk1 (DKD) knockdown on p130 and RB phosphorylation and expression of Ki67 and different cell cycle regulators in non-treated (NT) and U2OS cells exposed to bleomycin (Bleo) for 16 hours. Arrows show diminished Ki67 levels and pRb/p130 phosphorylation, accumulation of D-type cyclins and downregulation of Cdk6 upon Chk1 KD that are prevented by p21 knockdown. LC, loading control. **C.** Immunoblots showing effects of Chk1 (Ch1) or p21 knockdown on pocket protein phosphorylation and expression of different cell cycle regulators in non-treated (NT) and U2OS cells exposed to bleomycin (Bleo) for 7 days (7d). Black arrows: elevated p130 and pRb hyper-phosphorylation after p21 depletion. Red arrowhead: strong P^S317^-Chk1 signal in the absence of p21. Blue arrowhead: low Cdk6 levels. LC, loading control. **D.** Flow cytometry DNA content profiles of non-treated (NT) and U2OS cells exposed to bleomycin for 24 hr. Caffeine (Caf) and KU-55933 (Ku) were added one hour before treatment. **E.** Representative phase contrast images from a single video-microscopy sequence showing bleomycin-treated control (siCtl) and double Chk1/p21 knock down (siChk1/p21) U2OS cells at different times after exposure to the drug. Red rectangles show the inserts presented in Figure 6D. Loading controls (LC) were amido-black stained membranes. Black arrow: Chk1 KD; White arrow; p21 KD; asterisk: DKD

**Figure S9.**
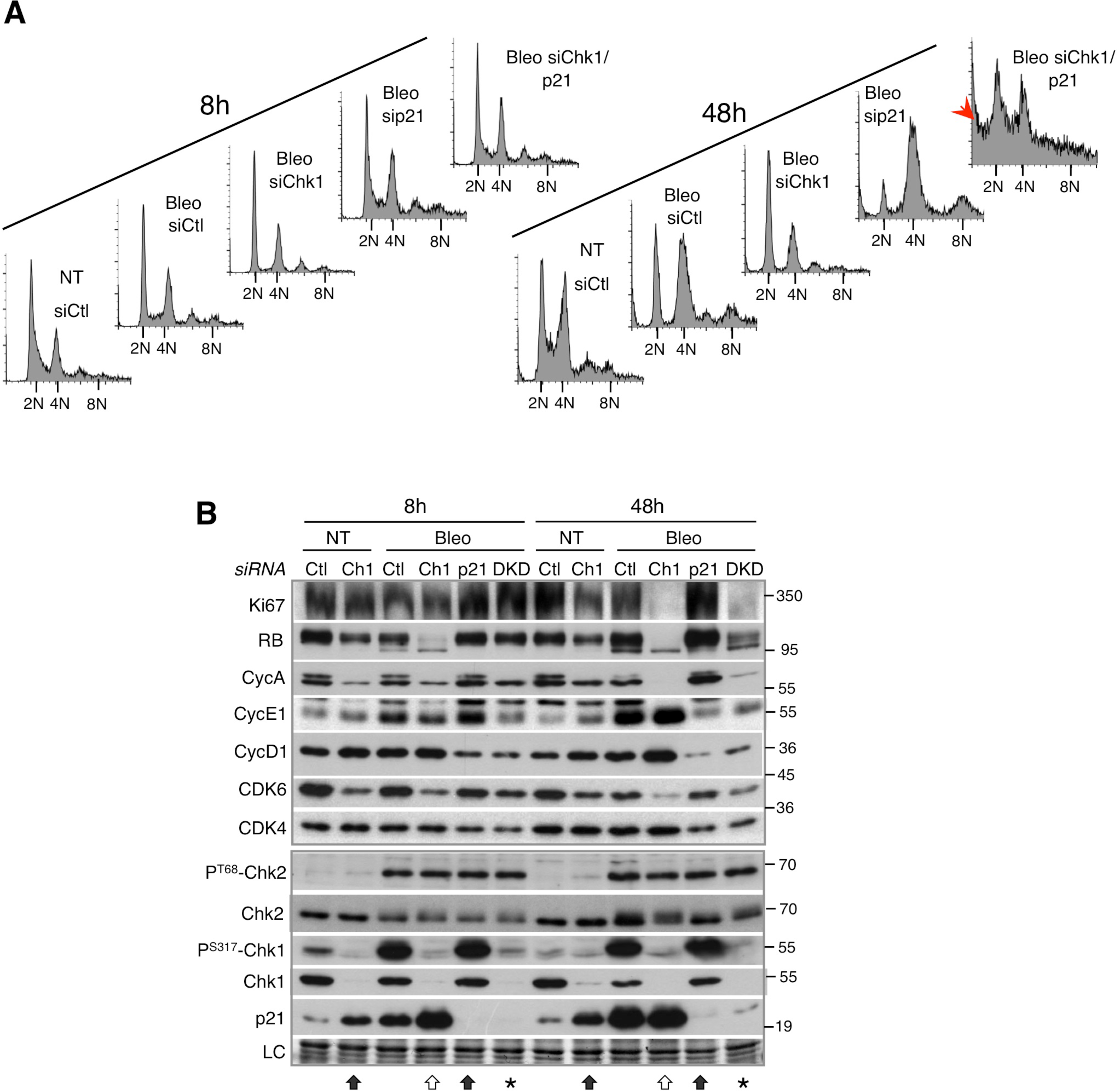
Accelerated DNA damage-induced cell cycle exit after Chk1 depletion in HCT-116 cells is mediated by p21 and CDK6 downregulation. **A.** Cell cycle analyses by flow cytometry DNA content of non-treated control (siCtl) and bleomycin-treated HCT-116 cells depleted for Chk1 (siCh1), p21 (sip21) or both (siChk1/21) proteins after 8 and 48 hours. Note the presence of sub-G1 population associated with dying cells after 48 hr in bleomycin-treated siChk1/21 cells (red arrow). **B.** Immunoblots showing effects of Chk1 (Ch1-black arrow), p21 (white arrow) or double p21/Chk1 (DKD-asterisk) knockdown on Ki67 levels, RB phosphorylation, expression of different cell cycle regulators and DNA damage response (P^T68^-Chk2, P^S317^-Chk1, p21) in non-treated (NT) and HCT-116 cells exposed to bleomycin (Bleo) for 8 and 48 hours. Loading controls (LC) are amido-black stained membranes.

